# Post-synaptic competition between calcineurin and PKA regulates mammalian sleep-wake cycles

**DOI:** 10.1101/2023.12.21.572751

**Authors:** Yimeng Wang, Siyu Cao, Daisuke Tone, Hiroshi Fujishima, Rikuhiro G. Yamada, Rei-ichiro Ohno, Shoi Shi, Kyoko Matsuzawa, Mari Kaneko, Maki Ukai-Tadenuma, Hideki Ukai, Carina Hanashima, Hiroshi Kiyonari, Kenta Sumiyama, Koji L. Ode, Hiroki R. Ueda

## Abstract

Phosphorylation of synaptic proteins is a pivotal biochemical reaction that controls the sleep-wake cycle in mammals. Protein phosphorylation in vivo is reversibly regulated by kinases and phosphatases. In this study, we investigated a pair of kinases and phosphatases that reciprocally regulate sleep duration. Through comprehensive screening of Protein kinase A (PKA) and phosphoprotein phosphatase (PPP) family genes via the generation of 40 gene knockout mouse lines including post-natal CRISPR targeting, we identified a regulatory subunit of PKA (*Prkar2b*), a regulatory subunit of protein phosphatase (PP) 1 (*Pppr1r9b*), and catalytic and regulatory subunits of PP2B (calcineurin) (*Ppp3ca* and *Ppp3r1*) as sleep control genes. AAV-mediated stimulation of PKA and PP1/calcineurin activities confirmed PKA as a wake-promoting kinase, while PP1 and calcineurin function as sleep-promoting phosphatases. The importance of these phosphatases in sleep regulation is supported by the dramatic changes in sleep duration associated with their increased and decreased activity, ranging from approximately 17.3 hours/day (PP1 expression) to 6.7 hours/day (post-natal CRISPR targeting of calcineurin). For these phosphatases to exert their sleep-promoting effects, localization signals to the excitatory post-synapse were necessary. Furthermore, the wake-promoting effect of PKA localized to the excitatory post-synapse negated the sleep-promoting effect of calcineurin, suggesting that PKA and calcineurin construct a hierarchical phosphorylation control network for sleep regulation at excitatory post-synapses.

## Main

Almost 75% of intracellular proteins undergo reversible phosphorylation modifications, dynamically regulating protein activities^1^. Reversible protein phosphorylation is also pivotal in the sleep-wake cycle regulation, a concept encapsulated as the “phosphorylation hypothesis of sleep regulation.^2^” Genetic screening studies have uncovered sleep-promoting kinases, such as CaMKIIα/β^3^, SIK3^4^, and ERK1/2^5^. Furthermore, mass-spectrometry-based phosphoproteomic analyses reveal that the phosphorylation state/level of synaptic proteins is profoundly influenced by sleep-wake states^6,7^. All such large-scale changes in phosphoprotein profiles are difficult to explain by a few sleep-promoting kinases: for example, proteins whose phosphorylation levels rise during sleep hint at the involvement of kinases in subsequent wakefulness induction. Moreover, the dynamic oscillation of phosphorylation levels throughout the sleep-wake cycle suggests that not only phosphorylation reactions but also the reverse reactions catalyzed by protein phosphatases play roles in sleep regulation.

Protein kinase A (PKA) is potentially involved in mammalian sleep control. PKA consists of catalytic and regulatory subunits. PKA’s catalytic subunit remains inactive when bound to the regulatory subunit. The regulatory subunit binds to cAMP, leading to dissociation from the catalytic subunit and PKA activation^8–10^. Studies in fruit flies suggest PKA activation promotes wakefulness through octopamine neurotransmitter release^11–15^. In addition, mammalian research indicates PKA may antagonize sleep induction via SIK1/2/3 phosphorylation^16,17^.

The presence of sleep-controlling kinases also points to the significance of protein phosphatases in the reversible control of phosphorylation residues during sleep regulation. However, the role of specific protein phosphatase activities in mammalian sleep regulation remains to be fully understood. Unlike kinases, only ∼10 catalytic subunits are encoded in the mammalian genome^18^, meaning phosphatase sequence specificity is generally low, making predictions of corresponding phosphatases from substrate sequences or specific kinases challenging. Regulatory subunits control a variety of functions, including inhibiting catalytic subunit activity and directing catalytic subunit localization^19^. Among the protein phosphatase family, Protein Phosphatase 2B (PP2B/calcineurin) has a unique Ca^2+^/CaM-activated regulatory mechanism^20^, and its role in synaptic plasticity regulation is well-documented^21,22^. Although fruit fly studies show calcineurin’s related influence on sleep duration^13,14^, it’s unclear whether calcineurin activation increases or decreases sleep. A comprehensive gene-knockout analysis could help decipher this complex interplay between catalytic and regulatory subunits of protein phosphatases in sleep regulation.

In our study, we found that PKA is the wake-promoting protein kinase in mammals, supported by both functional inhibition and functional enhancement perturbations. PKA appears to mainly exhibit its wake-promoting effects at the post-synapses of excitatory neurons. Given phosphorylation’s reversible nature in vivo, we also identified phosphatases that counteract the wake-promoting of PKA effect at excitatory post-synapses, discovering that Protein Phosphatase 1 (PP1) and calcineurin induce sleep selectively at excitatory post synapses. Based on these findings, we posit that the wake-promoting kinase PKA and sleep-promoting phosphatases (calcineurin and PP1) have opposing effects on sleep regulation at excitatory post-synapses.

### PKA is a wake-promoting kinase in mammals

To examine the role of each subunit of protein kinase A (PKA) in sleep regulation, we conducted a comprehensive gene knockout (KO) study (**Fig. 1a**). The mouse genome contains six genes in the PKA family: two are catalytic subunits (*Prkaca* and *Prkacb*), and four are regulatory subunits (*Prkar1a*, *Prkar1b*, *Prkar2a*, and *Prkar2b*). We employed the triple-target CRISPR method^23^, using three distinct guide RNAs to target a single gene, to knock out each member of the PKA family (**Extended Data Fig. 1a-e**). However, we excluded *Prkar1a* from our study since its KO mice are known to be lethal^24^. The genotypes of the KO mice were validated using qPCR (**Extended Data Fig. 1g-j**), and their phenotypes were assessed using a respiration-based sleep phenotyping system, termed the Snappy Sleep Stager (SSS) ^23^, under a 12:12 hr light-dark cycle. We found that *Prkaca* KO mice were also lethal, aligning with previous observations in the C57BL/6 background ^25^. Among the four viable KO mouse strains (*Prkacb*, *Prkar1b*, *Prkar2a*, and *Prkar2b* KO), we observed a significant reduction in sleep duration in the *Prkar2b* KO mice, averaging 583.4 ± 21.4 min (mean ± SEM, n = 8). This duration is approximately 141.6 min (or about 3.3 S.D.) less than that of the wild-type mice (p < 0.001) (**Fig. 1b**). The sleep/wake amplitude, defined as the coefficient of variation (CV: standard deviation divided by the mean) of sleep duration for each 10 min bin over 24 hours, was increased in the *Prkar2b* KO mice, likely due to decreased sleep during the dark phase (**Fig. 1b, c**). The reduced *P*_WS_ (transition probability from wakefulness to sleep) suggests that the wake phase was more stable in these mice (**Fig. 1b**). Using a different set of gRNA targets, we corroborated these sleep phenotypes, ruling out potential off-target effects from specific gRNA sequences (**Extended Data Fig. 1f, k**, and **2a, b**). Consistent with the observed extended wakefulness in the *Prkar2b* KO mice, these mice displayed an enhanced wakefulness response when introduced to a new environment, as seen in the cage change experiment during the dark phase (ZT = 12) (**Extended Data Fig. 2c, d**).

**Fig. 1.**
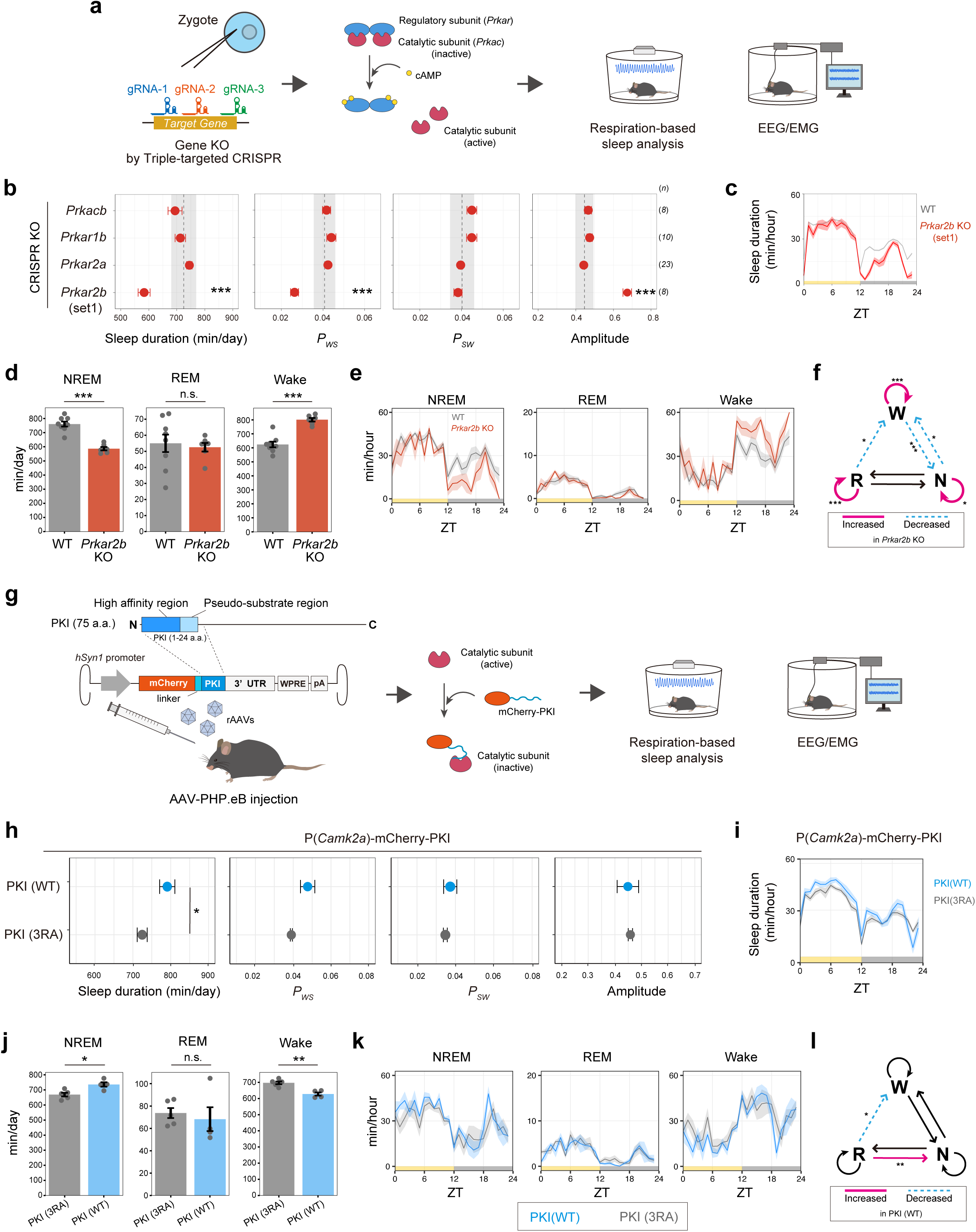
Identification of *Prkar2b* as a sleep controlling gene. **(a)** Schematic diagram of production of PKA KO mice by Triple-targeted CRISPR and sleep phenotyping. Genes encoding PKA catalytic subunit or regulatory subunit genes were targeted by the method. **(b, c)** Sleep/wake parameters (b) and sleep profiles (c) of the PKA KO mice, averaged over 6 days. Sleep duration is the total sleep duration in a day. Amplitude represents the variation of sleep duration per hour in a day. *P_WS_* and *P_SW_* are the transition probability between wakefulness and sleep. The black dashed line and the shaded area represent the mean and 1 SD range, respectively, of the wild-type mice wild-type C57BL/6N mice (n=108). The number of mice used in the analysis is shown as (*n*). Steel’s tests were performed between mutants and the wild-type C57BL/6N mice. **(d, e)** Sleep phenotypes (d) and sleep profiles (e) measured by EEG/EMG recordings for *Prkar2b* KO mice (n=6) and wild-type C57BL/6N mice (n=8). Student’s t-tests were performed between *Prkar2b* KO and the wild-type C57BL/6N mice. **(f)** Differences in transition probabilities (between wakefulness (W), NREM sleep (N), and REM sleep (R)) for *Prkar2b* KO mice. Magenta lines and dashed blue lines indicate when the values for *Prkar2b* KO mice are significantly (p < 0.05) higher and lower, respectively. Statistical analysis of *P_RW_* by Wilcoxon test and all others by Student’s t-test. **(g)** Schematic diagram of AAV-based PKI expression and sleep phenotyping. N-terminus region (1-24) of PKI were fused to mCherry and expressed in a mouse brain **(h, i)** Sleep parameters (h) and sleep profiles (i) of mice expressing PKI (WT) (n=6) and its inactive mutant (3RA) (n=6) under the *Camk2a* promoter, averaged over 6 days. The dosages of the AAVs were 4.0 x 10^11^ vg/mouse. Statistical analysis of sleep duration and *P_SW_* by Student’s t-test and *P_WS_* and Amplitude by Welch’s t-test. **(j, k)** Sleep phenotypes (j) and sleep profiles (k) measured by EEG/EMG recordings for PKI expressed mice (WT, n=4; 3RA, n=5). Student’s t-tests were performed. **(l)** Differences in transition probabilities (between wakefulness (W), NREM sleep (N), and REM sleep (R)) for PKI-expressing mice. Magenta lines and dashed blue lines indicate when the values for PKI (WT) mice are significantly (p < 0.05) higher and lower, respectively. Statistical analysis of *P_RW_* by Wilcoxon test, *P_NW_*, *P_NR,_ P_RN,_ P_RR_* by Student’s t-test, and all others by Welch’s t-test. Shaded areas in the line plots represent SEM. Error bars: SEM, *p < 0.05, **p < 0.01, ***p < 0.001. ZT, zeitgeber time; WT, wild-type.

To further investigate the sleep architecture of *Prkar2b* KO mice, we performed EEG/EMG recordings under a 12:12 hr light-dark cycle. These mice showed reduced NREM sleep duration, particularly during the dark phase (**Fig. 1d, e**). This reduction was primarily due to both a decreased *P*_WN_ (transition probability from wakefulness to NREM sleep) and an elevated *P*_WW_ (probability of remaining awake), reinforcing the idea of stabilized wakefulness in *Prkar2b* KO mice (**Fig. 1f**). The increased wakefulness coincided with a decrease in NREM delta power (**Extended Data Fig. 2e**), indicating PKA’s role in sleep pressure regulation.

Given that *Prkar2b* encodes a PKA regulatory subunit that inhibits the kinase activity of the PKA catalytic subunit, we next investigate if introducing a PKA inhibitor would induce the opposite sleep phenotype of *Prkar2b* KO mice. We expressed the PKA inhibitor peptide (PKI)^26,27^ in the mice brains, expressed under the *Camk2a* promoter using AAV.PHP.eB (**Fig. 1g**), a setup previously demonstrated to effectively induce gene expression throughout the mouse brain^28^. As, expected, PKI expression led to prolonged sleep duration when compared to its loss-of-function counterpart, PKI (3RA)^29^ (**Fig. 1h, i**). This observation further emphasizes the intrinsic role of PKA catalytic activity in promotingg wakefulness. EEG/EMG data also revealed that PKI-expressing mice experienced extended NREM sleep accompanied by an increased tendency toward NREM delta power (**Fig. 1j-l and Extended Data Fig. 2f**).

The effects of PKA inhibition could also be replicated by expressing the regulatory subunit. Expressing PKA regulatory subunit mutants that cannot bind cAMP is theorized to dampen the activation of endogenous PKA in a dominant-negative fashion (**Extended Data Fig. 3a**)^30,31^. As expected, introducing such mutants, *Prkar1a* (G325D) and *Prkar1b* (G325D), led to sleep increases akin to PKI expression (**Extended Data Fig. 3b-e**). These findings also suggest that the role of *Prkar1a* in sleep, unverified due to its embryonic lethality upon knockout^24^, aligns with that of *Prkar1b*. Although Prkar2a expression does not result in significant sleep change regardless of wild-type or dominant-negative mutant (**Extended Data Fig. 3f, g**), expression of *Prkar2b* showed a marked increase in sleep duration even in the case of wild-type *Prkar2a* expression (**Extended Data Fig. 3h, i**); note that expression of wildtype *Prkar2a* showed 807.3 min/day sleep while untreated mice show ∼750 min/day sleep in our assay system. This phenotype is opposite to the *Prkar2a* KO, thus supporting the importance of *Prkar2a* for sleep control.

### Cortical excitatory post-synapse is a candidate subcellular component for the wake-promoting function of PKA

We next examined whether PKA activation alone is sufficient to induce wakefulness. We expressed a constitutively active mutant of the PKA catalytic subunit, *Prkaca* (H88Q:W197R)^32^ (**Fig. 2a**). When compared to the expression of the wild-type *Prkaca*, the constitutive-active mutant *Prkaca* (H88Q:W197R) led to a reduction in sleep duration (**Fig. 2b**), underscoring PKA’s role as a wake-promoting kinase. This observation aligns with the sleep phenotype of *Prkaca* KO mice, where sleep alterations were primarily seen during the dark phase (**Fig. 2c**).

**Fig. 2.**
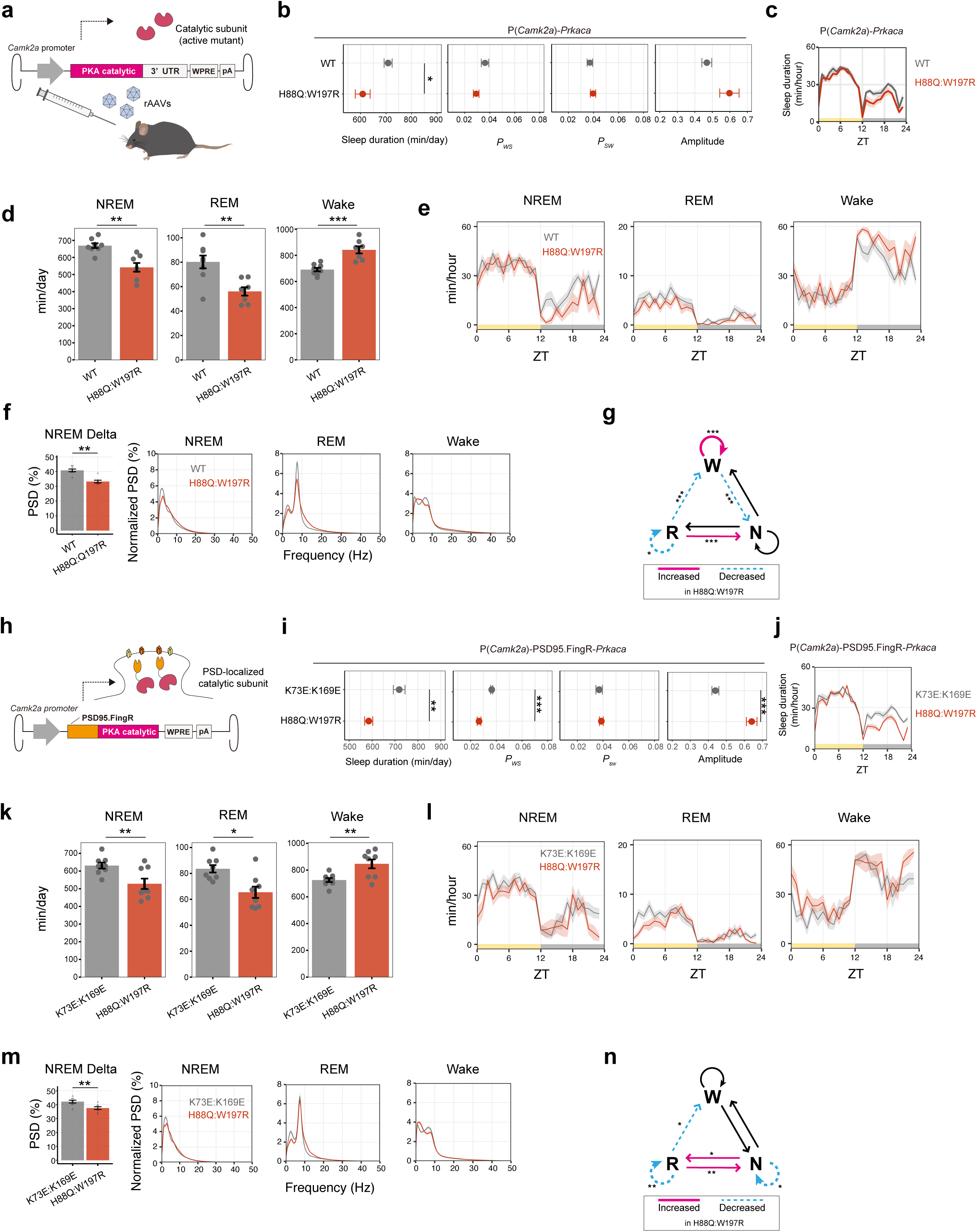
PKA is a sleep promoting kinase in mammals. **(a)** Schematic diagram of AAV-based PKA catalytic subunit expression. **(b, c)** Sleep/wake parameters (b) and sleep profiles (c) of mice expressing *Prkaca* WT (n=6) and its active mutant (H88Q:W197R) (n=6) under the *Camk2a* promoter, averaged over 6 days. The dosages of the AAVs were 1.0 x 10^10^ vg/mouse. Student’s t-tests were performed. **(d-e)** Sleep phenotypes (d) and sleep profiles (e) measured by EEG/EMG recordings for *Prkaca* (WT) (n=8) and *Prkaca* (H88Q:W197R) (n=7). Student’s t-tests were performed for comparisons. **(f)** NREM power density in delta domain (1-4 Hz) and EEG power spectra. Wilcoxon test was performed for the comparison in NREM delta power. **(g)** Differences in transition probabilities (between wakefulness (W), NREM sleep (N), and REM sleep (R)) between *Prkaca* (WT) and *Prkaca* (H88Q:W197R) mice under *Camk2a* promoter. Magenta lines and dashed blue lines indicate when the values for H88Q:W197R mice are significantly (p < 0.05) higher and lower. Student’s t-tests were performed for comparisons of the transition probabilities. **(h)** Schematic diagram of localized expression of PKA catalytic subunit based on PSD95.FingR. PSD95.FingR was fused to N-terminus of PKA catalytic subunit and expressed in mice brains. **(i, j)** Sleep/wake parameters (i) and sleep profiles (j) of mice expressing PSD95.FingR-*Prkaca* inactive mutant (K73E:K169E) (n=6) and its active mutant (H88Q:W197R) (n=6) under the *Camk2a* promoter. averaged over 6 days. The dosages of the AAVs were 5.0 x 10^9^ vg/mouse. Student’s t-tests were performed. **(k, l)** Sleep phenotypes (k) and sleep profiles (l) measured by EEG/EMG recordings for PSD95.FingR-*Prkaca* inactive mutant (K73E:K169E) (n=8) and its active mutant (H88Q:W197R) (n=8). Student’s t-tests were performed for comparisons. **(m)** NREM power density in delta domain (1-4 Hz) and EEG power spectra. Student’s t-test was performed for the comparison in NREM delta power. **(n)** Differences in transition probabilities (between wakefulness (W), NREM sleep (N), and REM sleep (R)) between PSD95.FingR-*Prkaca* inactive mutant (K73E:K169E) (n=8) and its active mutant (H88Q:W197R) (n=8). Magenta lines and dashed blue lines indicate when the values for H88Q:W197R mice are significantly (p < 0.05) higher and lower. Statistical analysis of *P_NN_* by Wilcoxon test, *P_RN_* by Welch’s t-test, and all others by Student’s t-test. Shaded areas in the line plots represent SEM. Error bars: SEM, *p < 0.05, **p < 0.01, ***p < 0.001. ZT, zeitgeber time; WT, wild-type.

EEG/EMG recordings from mice expressing *Prkaca* (H88Q:W197R) verified a notable reduction in NREM sleep, predominantly during the dark phase, accompanied by a decline in NREM delta power (**Fig. 2d-f**). Transition probabilities between NREM-REM-Wake states also showed that *Prkaca* (H88Q:W197R) expression led to an increase in *P*_WW_ and a decrease in *P*_WN_, indicating wake phase stabilization (**Fig. 2g**). The patterns of reduced NREM sleep in the dark phase and enhanced wakefulness are reminiscent of the *Prkar2b* KO mice phenotype. These results suggest that increased PKA kinase activity alone can stabilize and induce wakefulness. Notably, *Prkaca* (H88Q:W197R) expression also diminished REM sleep (**Fig. 2d**), implying that continuous wakefulness promotion by PKA can suppress REM sleep.

Previous phosphoproteomics research indicated that the sleep-wake cycle profoundly impacts the phosphorylation of synaptic proteins^6,7,33^. Given PKA’s established role in regulating synaptic plasticity at excitatory post-synapses^34^, we postulate that the excitatory post-synapse might be a pivotal subcellular component for the wake-promoting function of PKA. In support of this notion, fusion of the active *Prkaca* (H88Q:W197R) with the PSD95-binding FingR sequence^35,36^ (PSD95.FingR-*Prkaca*) remains the wake-promoting effect to the similar level with that of the non-localized expression of *Prkaca* (H88Q:W197R) (**Fig. 2h-j and Extended Data Fig. 4a-c**). For the control condition in these experiments, we utilized the loss-of-function *Prkaca* mutant K73E:K196E^37,38^. We also suggest wakefulness promotion upon expressing *Prkaca* (H88Q:W197R) mainly in cortical excitatory neurons, based on the wake-promoting effect of *Prkaca* (H88Q:W197R) expressed by the combination of the *Camk2a* promoter with Nex-Cre knock-in mice^39^ (**Extended Data Fig. 4d-f**).

Mice expressing excitatory post-synapse localized PSD95.FingR-*Prkaca* (H88Q:W197R) also exhibited reduced NREM and REM sleep and a decline in NREM delta power (**Fig. 2k-m**), though the wake stabilization was not statistically significant by the *P*_WW_ metric (**Fig. 2n**). In conclusion, we propose that PKA is a wake-promoting kinase and its wake-promoting influence operates at excitatory post-synapses.

### PP1 is a sleep-promoting phosphatase functioning at excitatory post-synapses

The phosphorylation status of proteins is reversibly regulated by kinases and protein phosphatases. Hence, we aimed to identify protein phosphatases that could counteract the wake-promoting function of PKA. The major Ser/Thr protein phosphatases expressed in the brain are PP1, PP2A, and calcineurin^18,40^. To investigate their roles, we undertook a comprehensive knockout (KO) experiment targeting these three phosphatase families (**Fig. 3a**). From the key genes associated with PP1, PP2A, and calcineurin, several are considered embryonically lethal or lead to substantial growth/developmental issues (*Ppp2r4*, *Strn*, *Strn3*, *Ppp2ca*, *Ppp2r1a*, *Ppp1cb*, *Ppp1r3c*, *Ppp1r8*, *Ppp1r10*, *Ppp1r12a*, *Trp53bp2*, *Ppp1r13l*, and *Ppp1r15b*) ^41,42^. A couple of genes were challenging for gRNA sequence selection due to an abundance of retroposons (*Ppp1cc* and *Ppp1r14b*). Consequently, we initiated a triple-target CRISPR-based screening for the genes depicted in **Extended Data Fig. 5**. KOs for *Ppp1r2*, *Ppp1r7, Ppp1r11*, *Ppp2r2a*, *Ppp2r3c*, *Ppp3ca*, and *Ppp3r1* were unattainable, suggesting an embryonically lethal phenotype. For the remaining 34 genes, we executed sleep recordings using the SSS-based system. We found that *Ppp1r9b* KO mice showed a pronounced sleep duration reduction and an enhanced sleep-wake rhythmicity amplitude (**Fig. 3b and Extended Data Fig. 6a, c**). These results were confirmed using a distinct set of triple-target CRISPR gRNA selections (**Extended Data Fig. 5** and 6b**, d, e**). Additionally, *Ppp1r3d* KO mice presented a minor yet statistically significant sleep duration increase (**Fig. 3b and Extended Data Fig. 6c**). Given the evident phenotype in *Ppp1r9b* KO mice, we then focused on understanding the sleep control mechanism of *Ppp1r9b*. EEG-EMG recording of *Ppp1r9b* KO mice revealed a significant reduction in both NREM and REM sleep (**Fig. 3c, d**), with a trend in decreasing NREM delta power (**Extended Data Fig. 6f**). Consistent with the increased amplitude observed in SSS recording, transition probabilities for the typical sequence from NREM, REM to awake states (*P*_WN_, *P*_NR_, and *P*_RW_) were all reduced in *Ppp1r9b* KO mice, indicating that both awake and NREM states are stabilized, as shown by the increased *P*_WW_ and *P*_NN_ (**Fig. 3e**).

**Fig. 3.**
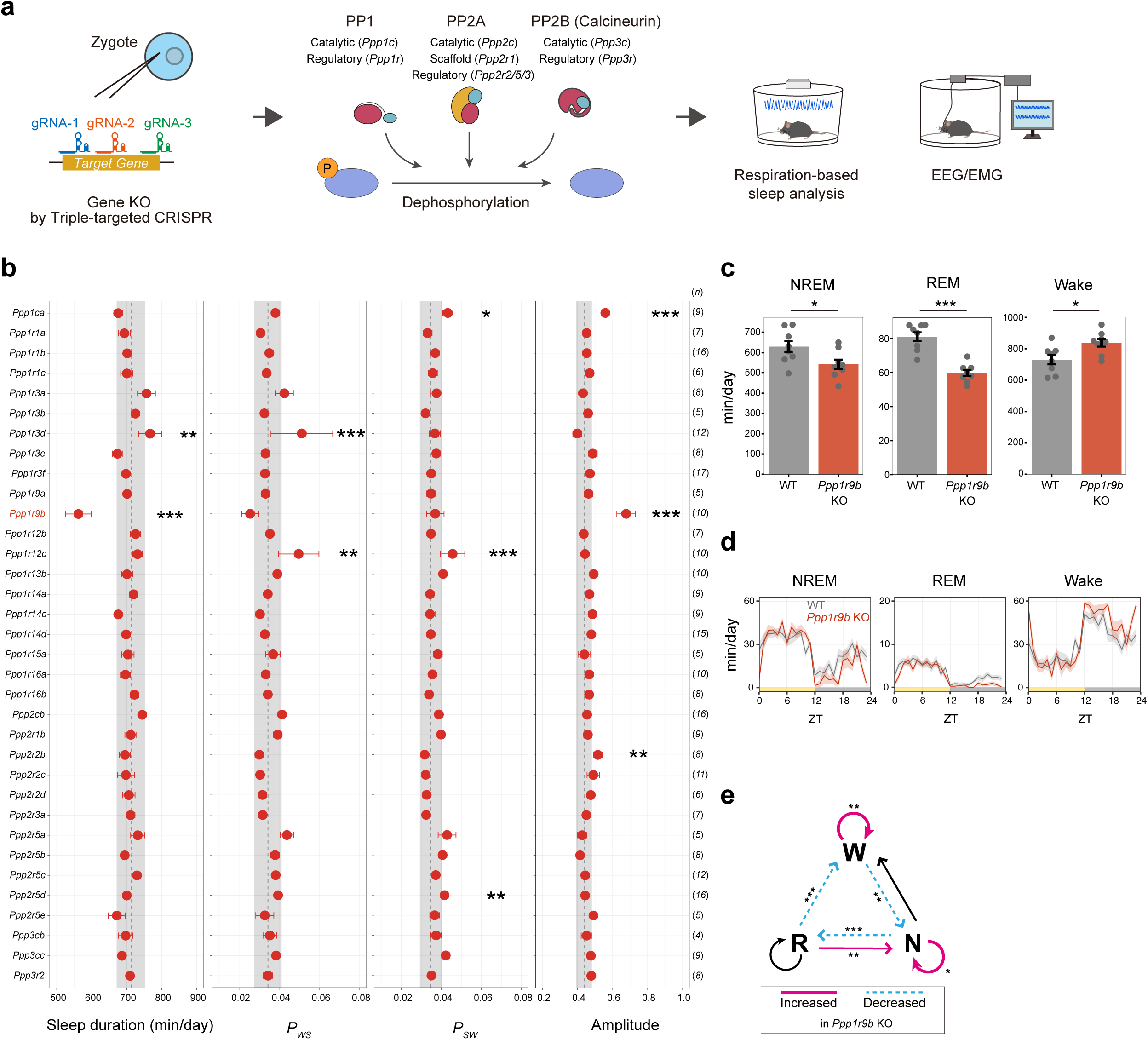
Identification of *Ppp1r9b* as a sleep controlling gene. **(a)** Schematic diagram of triple-target CRISPR method used for targeting PP1, PP2A, and PP2B (calcineurin) genes followed by sleep analyses using the respiration-based sleep phenotyping system (SSS) or EEG/EMG recording. Detailed information of designed gRNAs was provided in **Extended Data Fig. 5**. **(b)** Sleep parameters of each 8-weeks-old male KO mice. Sleep duration is the total sleep duration in a day. Amplitude represents the variation of sleep duration per hour in a day. *P_WS_* and *P_SW_* are the transition probability between wakefulness and sleep. The black dashed line and the shaded area represent the mean and 1 SD range, respectively, of the wild-type mice wild-type C57BL/6N mice (n=101). The number of mice used in the analysis is shown as (*n*). Dunnett’s tests were performed between mutants and the wild-type C57BL/6N mice. **(c, d)** Sleep phenotypes (c) and sleep profiles (d) measured by EEG/EMG recordings for *Ppp1r9b* KO mice (n=8) and wild-type C57BL/6N mice (n=8). Student’s t-tests were performed. **(j)** Differences in transition probabilities (between wakefulness (W), NREM sleep (N), and REM sleep (R)) between wild-type C57BL/6N mice and *Ppp1r9b* KO mice. Magenta lines and dashed blue lines indicate when the values for *Ppp1r9b* KO mice are significantly (p < 0.05) higher and lower. Statistical analysis of *P_RN_* by Welch’s t-test, and all others by Student’s t-test. Shaded areas in the line plots represent SEM. Error bars: SEM, *p < 0.05, **p < 0.01, ***p < 0.001. ZT, zeitgeber time; WT, wild-type.

*Ppp1r9b*, also known as Neurabin-2/Spinophilin, has been shown to target some PP1 catalytic subunits to excitatory post-synapses^43,44^. One anticipated result of *Ppp1r9b* KO is the absence of the PP1 catalytic subunit from these post-synapses. Accordingly, we examined whether reintroducing the PP1 catalytic subunit at excitatory post-synapses would counteract the *Ppp1r9b* KO effects (i.e., increased sleep and decreased amplitude) (**Fig. 4a**). Compared to the loss-of-function mutants of *PPP1CA* (H248K and D95N)^45,46^, AAV-based expression of wild-type *PPP1CA* and gain-of-function mutant lacking the inhibitory phosphorylation residue (T320A)^47^ resulted in increased sleep duration as well as decreased amplitude (**Fig. 4b, c**). Interestingly, such sleep effects were clearly observed only when *PPP1CA* was expressed as a fusion protein with PSD95; expression of *PPP1CA* without PSD95 fusion had a negligible effect on sleep duration (**Extended Data Fig. 7a-c**). However, amplitude reduction was still observed by the expression of wild-type or T320A *PPP1CA* without PSD95 fusion (**Extended Data Fig. 7b, c**). Interestingly, the KO of *Ppp1ca* showed an amplitude increase without showing a strong effect on sleep duration (**Fig. 3b**). These results suggest that *PPP1CA* (and *PPP1R9B*) have distinct NREM-sleep-promoting functions mainly at the excitatory post-synapses and sleep-wake transition control involving non-post-synapse regions.

**Fig. 4.**
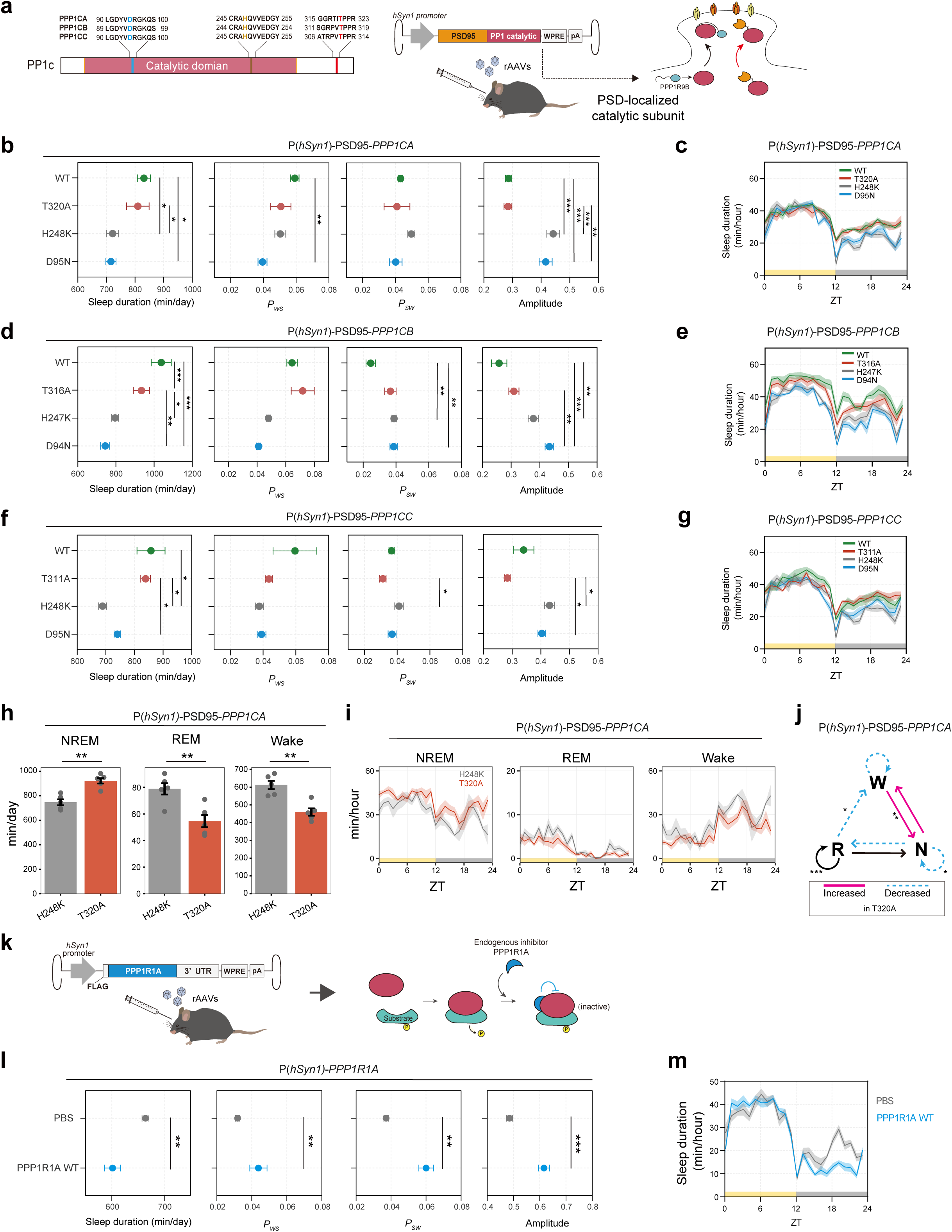
PP1 is a sleep promoting phosphatase at excitatory post-synapses. **(a)** Information of the point mutations on PP1c genes used in this study (left). Red-, dark yellow-, blue-colored sites indicate mutations of constitutive active, activity-dead, dominant negative, respectively. The Schematic diagram of AAV-PHP.eB based expression of PSD95-fused PP1 catalytic subunits (*PPP1CA*, *PPP1CB*, *PPP1CC*) in mice postsynaptic density (right). PSD95 fusion emulated synaptic-translocation of PPP1R9B-PP1c complex. AAVs were injected into 6-weeks-old male wild-type C57BL/6N mice with the dosage of 4×10^11^ vg/mouse. **(b-g)** Sleep parameters (b, d, f) and 24-hour sleep profile (c, e, g) of PSD95-fused PP1c-expressing mice (n=6 each) measured by SSS, averaged over 6 days. For *PPP1CA*, Tukey-Kramer tests were performed for sleep duration, *P_WS_* and amplitude, and Steel-Dwass was performed for *P_SW_* between all individual groups. For *PPP1CB*, Tukey-Kramer tests were performed for sleep duration, *P_SW_* and amplitude, and Steel-Dwass test was performed for *P_WS_*. For *PPP1CC*, Tukey-Kramer tests were performed for sleep duration and amplitude, and Steel-Dwass test was performed for *P_WS_* and *P_SW_*. **(h, i)** Sleep phenotypes (h) and sleep profiles (i) measured by EEG/EMG recordings for PSD95-fused *PPP1CA* (H248K) (n=5) and T320A mutant mice (n=5). Student’s t-tests were performed for the comparisons. The dosages of the AAVs were 1.5 x 10^11^ vg/mouse. **(j)** Differences in transition probabilities (between wakefulness (W), NREM sleep (N), and REM sleep (R)) between mice expressing PSD95-fused *PPP1CA* H248K or T320A mutant. Magenta lines and dashed blue lines indicate when the values for T320A mice are significantly (p < 0.05) higher and lower. Statistical analysis of *P_NW_*, *P_WN_*, *and P_WW_* by Welch’s t-test, and all others by Student’s t-test. **(k)** Schematic diagram of expression of endogenous PP1 inhibitor *PPP1R1A* gene with AAV-PHP.eB. **(l, m)** Sleep parameters (l) and 24 hour sleep profile (m) of 8-weeks-old PBS injected mice (n=6) and *PPP1R1A*-expressing mice (n=6) measured by SSS, averaged over 6 days. AAVs were injected into 6-weeks-old male wild-type C57BL/6N mice with the dosage of 4×10^11^ vg/mouse. Student’s t-test were performed. Shaded areas in the line plots represent SEM. Error bars: SEM, *p < 0.05, **p < 0.01, ***p < 0.001. ZT, zeitgeber time; WT, wild-type.

*PPP1CB* and *PPP1CC* also required PSD95 fusion to exert a clear increase in sleep duration (**Fig. 4d-g and Extended Data Fig. 7d-g**), suggesting that excitatory post-synapses are the central location for the sleep-promoting function of PP1 enzymatic activity. The sleep-promoting function of PP1 is clearly attributed to the increase of NREM sleep compared with the loss-of-function H248K (**Fig. 4h, i**), though the change in the NREM delta power was not significant (**Extended Data Fig. 8a**). REM sleep amount is even reduced in the case of gain-of-function PSD95-*PPP1CA* (T320A) (**Fig. 4h**). Consistent with the decreased amplitude under the expression of PSD95-*PPP1CA* (T320A) (**Fig. 4b**), both wake stability (*P*_WW_) and NREM sleep stability (*P*_NN_) were significantly reduced (**Fig. 4j**).

The role of the endogenous PP1 catalytic subunit for sleep promotion was further confirmed by the decreased sleep duration under the expression of *PPP1R1A*, a regulatory subunit that inhibits the PP1 catalytic subunit^19^ (**Fig. 4k-m**). Overall, our results characterize PP1 as a sleep-promoting phosphatase functioning at the excitatory post-synapses. The sleep-promoting function is still evident under forced awakening conditions: *Ppp1r9b* KO mice showed enhanced awake duration induction by cage change (**Extended Data Fig. 8b**). On the other hand, sleep-promoted PSD95-*PPP1CA* (T320A)) or PSD95-*PPP1CC* (T311A) mice showed reduced awake duration induction by cage change (**Extended Data Fig. 8c, e**), while the effect of PSD95-*PPP1CB* (T316A) was not evident (**Extended Data Fig. 8d**).

### Calcineurin is a potent NREM-sleep-promoting phosphatase functioning at excitatory post-synapses

Whether calcineurin serves to promote sleep or wakefulness is still unclear in studies in flies^13,14^. In our KO screening, viable calcineurin subunit KO mice (i.e., *Ppp3cb*, *Ppp3cc*, and *Ppp3r2* KO) did not show clear sleep phenotypes. Thus, we decided to investigate the role of *Ppp3ca* and *Ppp3r1*, which would otherwise result in embryonic lethality in the KO mice, by implementing post-natal KO via AAV-mediated expression of triple-target CRISPR^48^ (**Fig. 5a**).

**Fig. 5.**
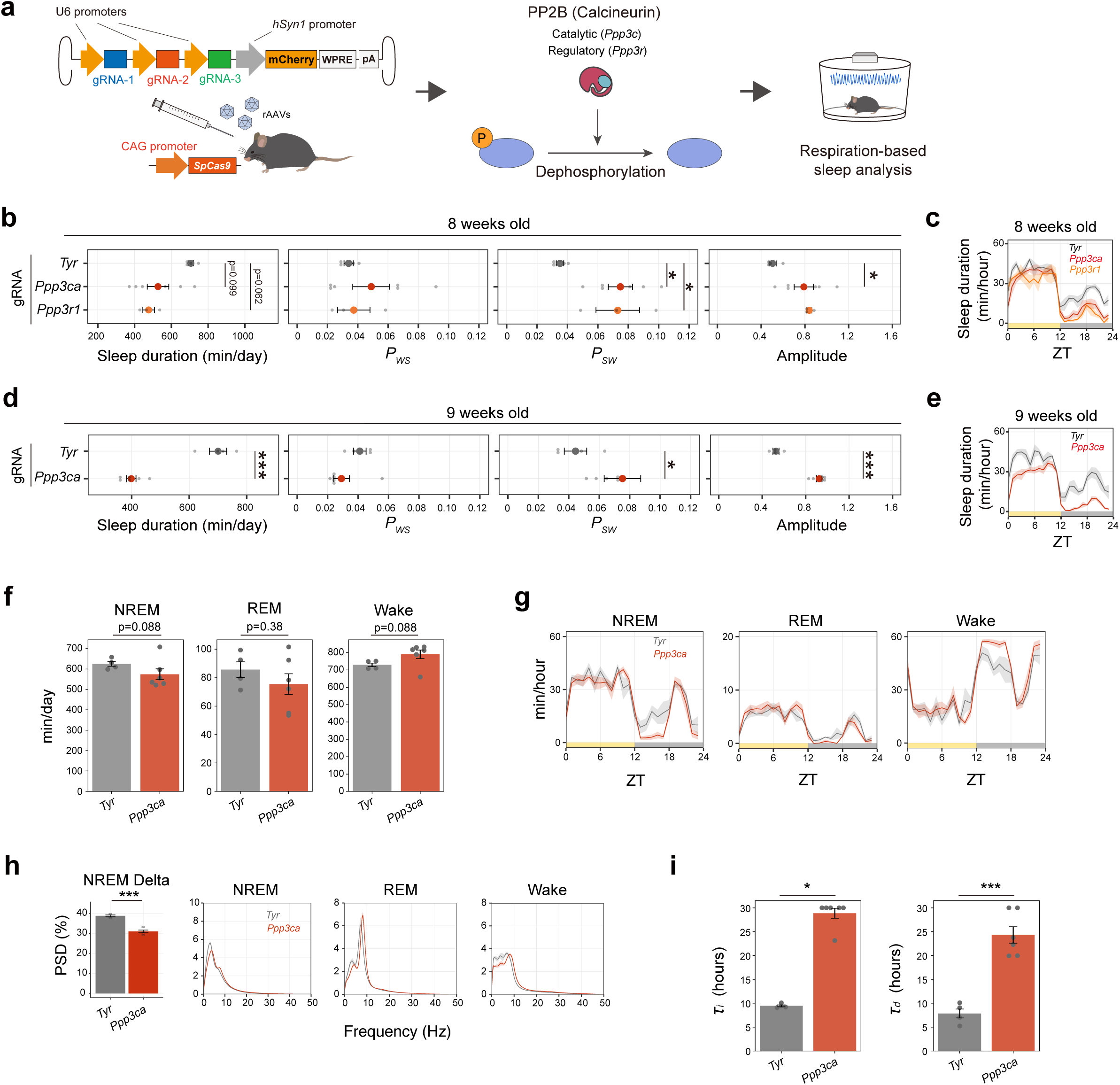
Calcineurin is critical for quantitative and qualitative control of sleep. **(a)** Schematic diagram of post-natal CRISPR KO method used for targeting PP2B catalytic and regulatory genes with H11-Cas9 knock-in mice, followed by the respiration-based sleep phenotyping system (SSS). **(b-e)** Sleep/wake parameters (b, d) and sleep profiles (c, e) of Tyrosinase (*Tyr*) (n=4), *Ppp3ca* (n=6) and *Ppp3r1* (n=6) -targeted mice at 8-week-old (b, c) and 9-week-old (d, e), averaged over 6 days. For 8-week-old mice, statistical analysis of sleep duration and amplitude by Steel’s test, and *P_WS_* and *P_SW_* by Dunnett’s test against *Tyr* KO group. For 9-week-old mice, statistical analysis of sleep duration and amplitude by Student’s t-test, and *P_WS_* and *P_SW_* by Wilcoxon test between *Tyr*- and *Ppp3ca*-tageted mice. **(f, g)** Sleep phenotypes (f) and sleep profiles (g) measured by EEG/EMG recordings for *Tyr* (n=4) and *Ppp3ca* (n=6) -targeted mice. Student’s t-test was performed for comparison of REM duration, Wilcoxon tests for NREM and wake durations. **(h)** NREM power density in delta domain (1-4 Hz) and EEG power spectra of *Tyr* and *Ppp3ca* -targeted mice. Student’s t-test was performed for the comparison in NREM delta power. **(i)** Estimated time constants for the increase of EEG delta power during awake/REM periods (τi) and the decrease of NREM EEG delta power during the NREM period (τd). Shaded areas in the line plots represent SEM. Error bars: SEM, *p < 0.05, **p < 0.01, ***p < 0.001. KO, knock out; ZT, zeitgeber time.

Reduced sleep duration was observed in Cas9-expressing mice with the administration of AAVs expressing triple CRISPR gRNAs targeting either *Ppp3ca* or *Ppp3r1*, compared to the control AAV expressing gRNAs targeting *Tyrosinase* (**Fig. 5b, c**). A more pronounced reduction in sleep duration for *Ppp3ca*-targeted mice was observed in the recordings from 9-week-old mice (**Fig. 5d, e**). This data suggests that in mammals, calcineurin may function as a sleep-promoting protein phosphatase. The reduced NREM sleep amount in *Ppp3ca* KO mice was confirmed by EEG-EMG recording (**Fig. 5f, g**). We also observed reduced NREM delta power in the KO mice (**Fig. 5h**). Furthermore, *Ppp3ca* KO mice had sever defect in the increase/decrease dynamics of NREM delta power. Estimation of increase and decrease time constant of delta power during awake and NREM epochs^49^ revealed that both increases and decrease rates are significantly slower in the *Ppp3ca* KO mice (**Fig. 5i**). These dramatic changes in sleep duration and NREM delta power dynamics suggest that calcineurin is a central regulator of sleep.

To further verify this sleep-promoting effect, we expressed a constitutively active variant of the calcineurin catalytic subunit. The catalytic domain of calcineurin is inhibited by its C-terminal domain. Therefore, deleting the C-terminal domain results in a constitutively active calcineurin mutant that operates independently from Ca^2+^/CaM activation^50,51^ (**Fig. 6a**). We found that the expression of constitutively active *PPP3CA* or *PPP3CB* similarly promotes sleep compared to both the wild-type and the phosphatase-inactive version (H151/160Q)^51,52^ of the C-terminal deleted forms (**Fig. 6b-e**). Notably, the sleep-promotion effect requires fusion with PSD95.FingR, and expressing *PPP3CA* and *PPP3CB* without this fusion did not significantly alter the sleep phenotype (**Extended Data Fig. 9a-e**). On the other hand, PSD95.FingR-fused *PPP3CA* and *PPP3CB* induced sleep increase even with a lower dosage of AAV, indicating the robust sleep-promoting effect of calcineurin at the excitatory post-synapses (**Extended Data Fig. 9f-j**). As in the case of PP1, the sleep-promoting effect of calcineurin can be observed in the cage change condition (**Extended Data Fig. 10a-d**). Specifically, the sleep-promoting PSD95.FingR-*PPP3CA* (1-389) or *PPP3CB* (1-408) mitigated the wakefulness elicited by exposure to a new environment, especially during the light phase (ZT = 0).

**Fig. 6.**
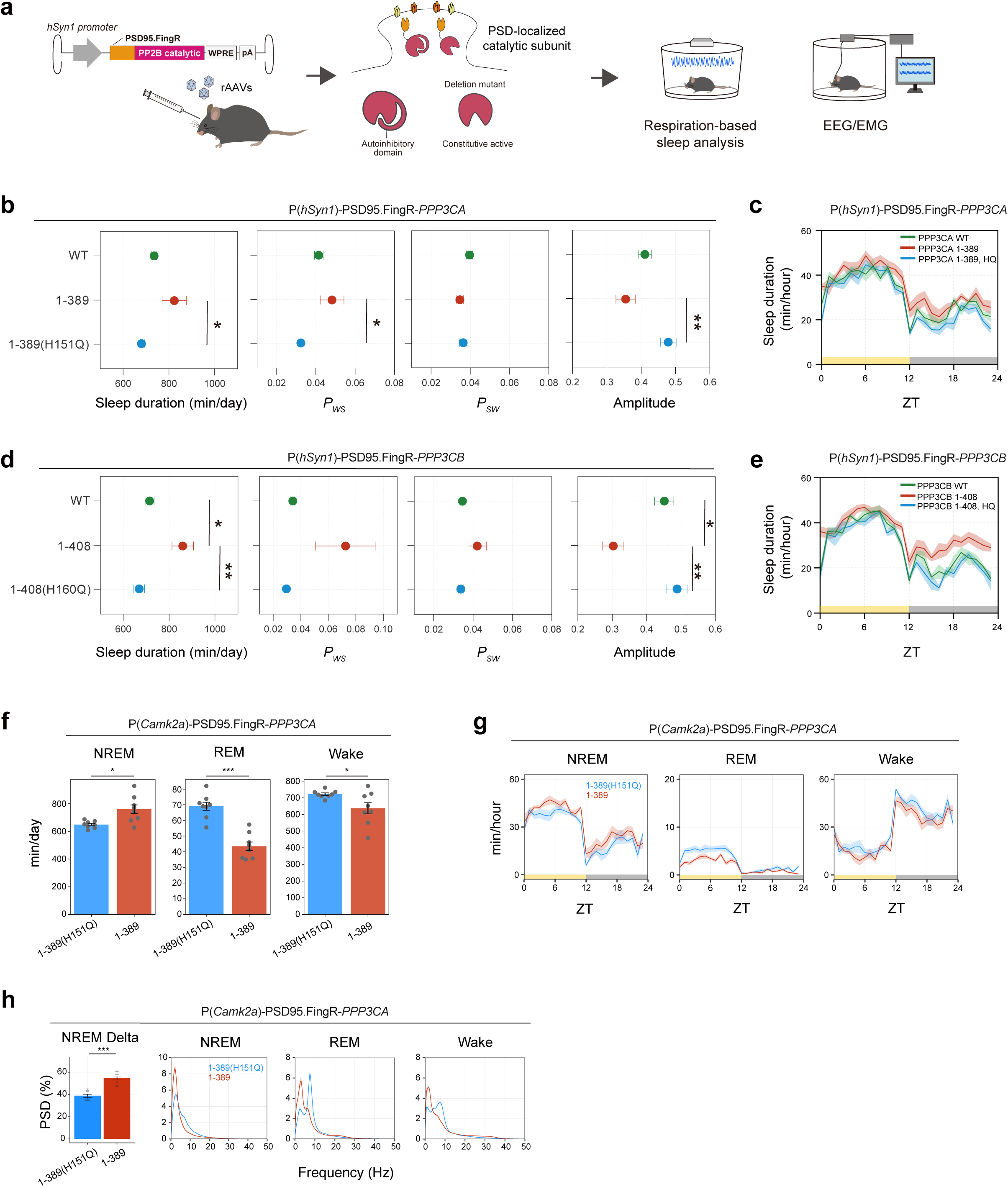
Calcineurin is a NREM sleep promoting phosphatase at excitatory post-synapses. **(a)** Schematic diagram of AAV-PHP.eB based expression of PSD95.FingR-fused PP2B catalytic subunits (PPP3CA, PPP3CB) in postsynaptic density. The catalytic domain in the PP2B catalytic subunit is kept blocked by its autoinhibitory domain under a low calcium concentration, while its activity will be released from the deletion of the autoinhibitory domain (e.g., constitutive active). The dosages of the AAVs were 5×10^10^ vg/mouse. **(b-e)** Sleep/wake parameters (b, d) and sleep profiles (c, e) of mice expressing PSD95.FingR-PP2B catalytic subunits, averaged over 6 days. WT, wild-type (n=6); 1-389, constitutive active deletion mutant (n=6); 1-389(H151Q), deletion mutant with inactive mutation (n=6). For PPP3CA, Steel-Dwass tests were performed for sleep duration and *P_WS_*, and Tukey-Kramer was performed for *P_SW_* and amplitude between all individual groups. For PPP3CB, Tukey-Kramer tests were performed for sleep duration, *P_WS_* and amplitude, and Steel-Dwass was performed for *P_SW_*. **(f, g)** Sleep phenotypes (f) and sleep profiles (g) measured by EEG/EMG recordings for mice expressing PSD95.FingR-*PPP3CA* under *Camk2a* promoter. 1-389, constitutive active deletion mutant (n=8); 1-389(H151Q), deletion mutant with inactive mutation (n=8). Student’s t-test was performed for comparison of REM duration, Welch’s t-tests for NREM and wake durations. **(h)** NREM power density in delta domain (1-4 Hz) and EEG power spectra of mice expressing PSD95.FingR-*PPP3CA* under *Camk2a* promoter. Student’s t-test was performed for the comparison in NREM delta power. Shaded areas in the line plots represent SEM. Error bars: SEM, *p < 0.05, **p < 0.01, ***p < 0.001. ZT, zeitgeber time; WT, wild-type.

The enhanced sleep duration can be attributed to the increased NREM sleep based on the EEG/EMG recording (**Extended Data Fig. 10e-f**). The NREM delta power profile is not significantly altered in the PSD95.FingR-*PPP3CA* (1-389) expressed under the control of *Syn1* promoter (**Extended Data Fig. 10g**). Interestingly, drastic increase of NREM delta power is observed focused expression of PSD95.FingR-*PPP3CA* (1-389) in the excitatory neurons by using *Camk2a* promoter (**Fig. 6f-h**), further confirming the role of calcineurin in the induction of NREM sleep and NREM delta power operating primarily at the excitatory post-synapse.

### Postsynaptic competition between PKA and calcineurin

Our results support the idea that wake-promoting PKA and sleep-promoting calcineurin both regulate sleep at the excitatory post-synapse. To decipher how these antagonistic kinases and phosphatases collaboratively influence overall sleep duration, we conducted a co-activation experiment involving both PKA and calcineurin (**Fig. 7a**). For the study, we expressed either or both of the constitutively active forms of PKA and calcineurin as PSD95.FingR fusion proteins. In the presence of loss-of-function PKA (PSD95.FingR-*Prkaca* (K73E:K169E)), constitutively active PSD95.FingR-*PPP3CA* (1-389) showed enhanced sleep duration compared to the co-expression of loss-of-function PSD95.FingR-*PPP3CA* (1-389:H151Q) (**Fig. 7b-c**). The enhanced sleep duration in the presence of constitutively active calcineurin is still evident in the co-presence of constitutively active PKA (PSD95.FingR-*Prkaca* (H88Q:W197R)). However, the absolute sleep duration is reduced in this condition when compared to the corresponding calcineurin conditions in the presence of loss-of-function PKA (PSD95.FingR-*Prkaca* (K73E:K169E)). This result suggests that both of sleep-inducing effect of calcineurin and wake-inducing effect of PKA are detectable in this co-expression setting and the total sleep duration is determined by the summation of the competing sleep/wake control effect of calcineurin and PKA.

**Fig. 7.**
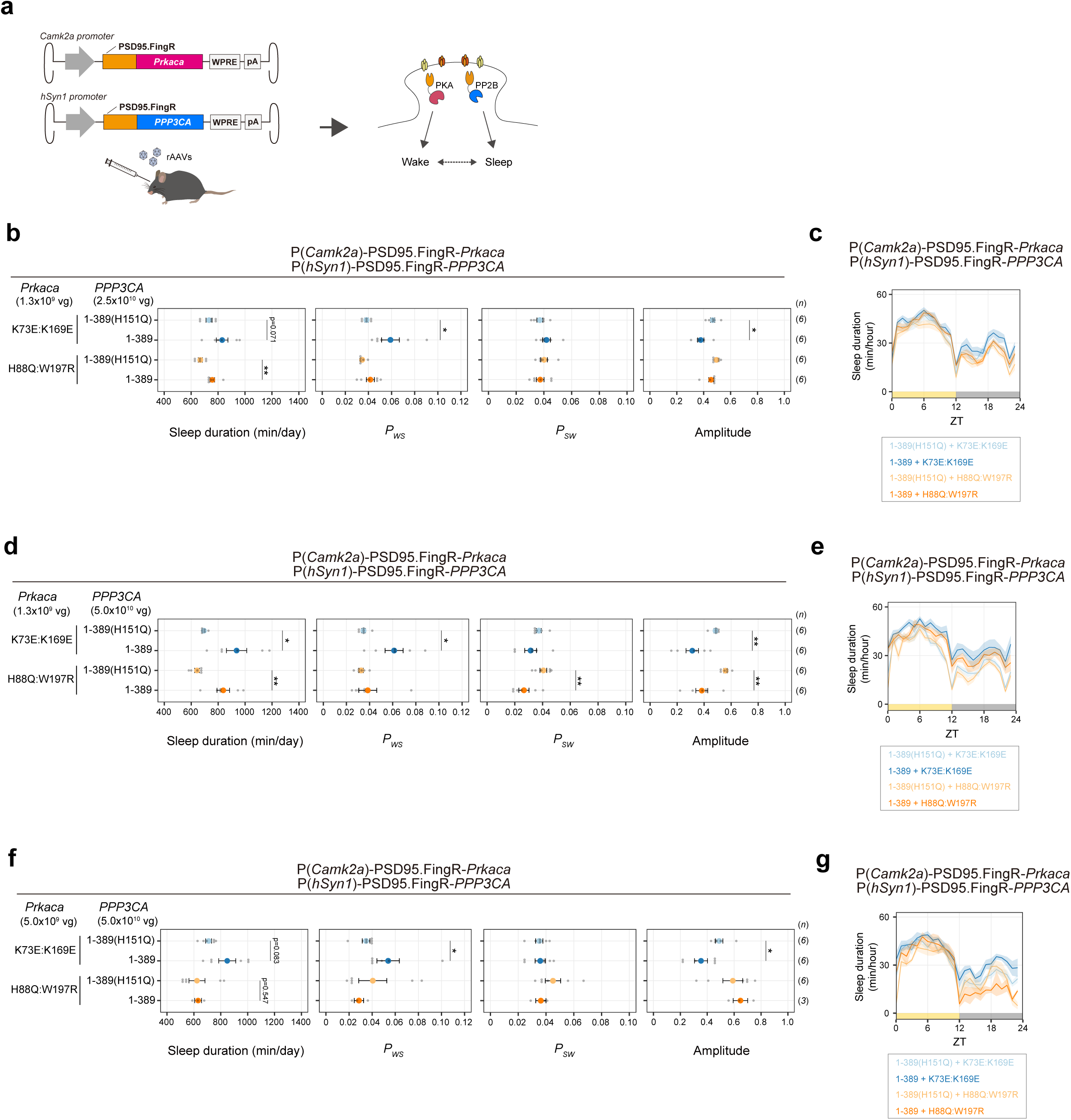
PKA competes with calcineurin at excitatory post-synapses in sleep control. **(a)** Schematic diagram of co-expression analysis of PSD95.FingR-*Prkaca* and PSD95.FingR-*PPP3CA*. The functional relationship between PKA as a sleep-promoting factor and calcineurin as a wake-promoting factor within postsynaptic density was investigated. **(b-g)** Sleep/wake parameters (b, d, f) and sleep profiles (c, e, g) of the PKA: calcineurin co-expressing mice, averaged over 6 days. Sleep duration is the total sleep duration in a day. Amplitude represents the variation of sleep duration per hour in a day. *P_WS_* and *P_SW_* are the transition probability between wakefulness and sleep. AAV dosages for *Prkaca* were 1.3×10^9^ (lower) (b-e) and 5.0×10^9^ (higher) (f, g). *Prkaca* KE:KE, kinase-dead; HQ:WR, constitutive active. AAV dosages for *PPP3CA* were 2.5×10^10^ (lower) (b, c) and 5.0×10^10^ (Higher) (d-g). *PPP3CA* 1-389(H151Q), inactive; 1-389, constitutive active. The number of mice used in the analysis is shown as (*n*). Statistical analyses were performed between 1-389 (H151Q) and 1-389 mice. In PKA (lower): Calcineurin (lower) condition (b), Under *Prkaca* (K73E:K169E) -expressing condition, statistical analysis of sleep duration, *P_SW_*, and amplitude by Student’s t-test, *P_WS_* by Welch’s t-test. Under *Prkaca* (H88Q:W197R) -expressing condition, statistical analysis of sleep duration and amplitude by Wilcoxon test, *P_WS_* by Welch’s t-test, and *P_SW_* by Student’s t-test. In PKA (lower): Calcineurin (higher) condition (d), Under *Prkaca* (K73E:K169E) - expressing condition, statistical analysis of sleep duration and *P_WS_* by Welch’s t-test, *P_SW_* and amplitude by Wilcoxon test. Under *Prkaca* (H88Q:W197R) -expressing condition, statistical analysis of sleep duration and amplitude by Welch’s t-test, *P_WS_* by Wilcoxon test, and *P_SW_* by Student’s t-test. In PKA (higher): Calcineurin (higher) condition (f), Under *Prkaca* (K73E:K169E) -expressing condition, statistical analysis of sleep duration by Welch’s t-test, *P_WS_* by Wilcoxon test, and *P_SW_* and amplitude by Student’s t-test. Under *Prkaca* (H88Q:W197R) -expressing condition, statistical analysis of sleep duration and *P_WS_* by Wilcoxon test, and *P_SW_* and amplitude by Student’s t-test. Shaded areas in the line plots represent SEM. Error bars: SEM, *p < 0.05, **p < 0.01, ***p < 0.001. ZT, zeitgeber time.

The additive effect of sleep-promoting calcineurin and wake-promoting PKA at the excitatory post-synapse is confirmed by changing the relative dosage of AAV for each enzyme. **Fig. 6d** and **6e** showed the results of the experiment similar to the **Fig. 6b** except for that the higher dosage of AAV expressing calcineurin. In this condition, sleep promotion effect of constitutively active calcineurin became more evident. On the other hand, increased sleep duration elicited by the constitutive active calcineurin was not observed under higher dosage of constitutively active PKA (PSD95.FingR-*Prkaca* (H88Q:W197R)) (**Fig. 6f, g**). In other words, the sleep promotion mediated by calcineurin is nullified by the higher level of the constitutive-active PKA. Thus, this co-expression study illuminates a competing relationship between the awake-promotion by PKA and sleep-promotion by calcineurin in governing sleep at the excitatory post-synapse.

## Discussion

Recent genetic researches have identified sleep-promoting kinases^3–5^. On the other hand, phosphoproteomics studies indicate that, while there is a general trend of increased phosphorylation levels of synaptic proteins during sleep deprivation^33^, there are not only phosphorylation sites that increase during wakefulness but also numerous sites that increase during sleep^6,7^. This makes it challenging to explain the phosphorylation state of synaptic proteins in sleep-wake regulation solely from the activity of sleep-promoting kinases. In this paper, we identified PKA as a wake-promoting kinase and PP1 and calcineurin as sleep-promoting phosphatases. The necessity and sufficiency of each kinase/phosphatase in the sleep duration determination process in mammals are supported by the observation that opposing changes in sleep duration occur with inhibition and activation of each enzymatic activity (for example, PKA; inhibition by PKI and activation by *Prkaca* H88Q:W197R, PP1; inhibition by PPP1R1A and activation by the expression of PP1 catalytic subunits, calcineurin: inhibition by post-natal CRISPR and activation by the expression of calcineurin catalytic subunits). These are positioned as having complementary roles to the sleep-promoting kinases such as CaMKIIα/β, MAPK, SIK3.

The relationship between PKA and calcineurin with sleep regulation has been demonstrated in studies using fruit flies^11–15^. The identified role of PKA as a wake-promoting kinase is consistent with a report demonstrating that the induced expression of PKA in mushroom bodies, which are thought to be a sleep center of *Drosophila*, decreases the duration of sleep^53^. A recent proteomics approach which investigated proteins expressed in the postsynaptic density of the mouse cortex and hippocampus during either wakefulness or sleep revealed that PKA components including both catalytic and regulatory subunits are enriched during the wake phase^6^. Furthermore, motif analysis of phosphopeptide predicted that PKA activity is enriched both in wake phase and sleep phase^6^, suggesting that PKA activity is dynamically controlled and substrate preference may be altered during sleep-wake cycle. The downstream factors involved in PKA’s wake regulation remain a subject for future research. Introduction of mutations to the conserved PKA phosphorylation sites in SIK1/2/3 results in sleep promotion^16,17^; thus, the SIK3-HDAC4 pathway^54,55^ might be a potential target of the awake-promoting function of PKA. However, because the phosphorylation mimic mutations at the PKA target sites in SIK3 also induce sleep^16^, further verification is needed to determine the molecular targets of the wake-promoting effects of PKA.

The role of calcineurin in sleep regulation in fruit flies has been ambiguous, as both genetic interventions that increased and decreased calcineurin activity led to diminished sleep^14^. However, in our mammalian study, we observed contrasting sleep phenotypes in relation to calcineurin activity: sleep duration decreased following post-natal CRISPR targeting of *Ppp3ca* and *Ppp3r1*, whereas it increased upon expression of *PPP3CA*/*PPP3CB* at the excitatory synapse. Consequently, in mammals, the primary role of calcineurin in sleep regulation appears to be in promoting sleep. Further research is necessary to investigate the downstream targets of calcineurin in sleep regulation, as well as the factors controlling calcineurin activity itself. It has been reported that the expression of calcineurin increases during sleep^56^, suggesting that calcineurin activity might be dynamically regulated throughout the sleep-wake cycle. We have proposed the significance of calcium signaling in sleep regulation, primarily focusing on CaMKII^3,57^: if calcineurin as well as CaMKII promote sleep downstream of calcium signaling, it should be interesting to investigate how CaMKII and calcineurin might divide their sleep-promoting roles. Given that CaMKII and calcineurin have different response characteristics to calcium signaling^58^, each enzyme might induce sleep in response to distinct types of calcium signals.

The relationship between sleep regulation and synaptic plasticity or synaptic homeostasis is well-established^59,60^. The enzymes PKA, PP1, and CaN, which were highlighted in our recent study, are also known for their roles in synaptic plasticity^61^. PKA is recognized as a factor responsible for memory consolidation including the consolidation during sleep^62^. Neurabin-2 has been implicated in neural plasticity and memory consolidation as well^63–65^, while the enzymatic activity of PP1 is also involved in memory extinction^66^. In addition, inhibition or overexpression of calcineurin showed abnormalities in memory consolidation and synaptic plasticity^63–65^, and thus calcineurin is understood to be one of the central factors in the control of synaptic plasticity^67^. A recent study in flies also suggests the involvement of IP_3_R both in sleep control and synaptic downscaling as a downstream factor of calcineurin^68^. Our observation about the catalytic subunits of PP1 and calcineurin in sleep regulation clearly requiring localization to the post-synapse indicates that these phosphatases function at the post-synapse for sleep control. The co-expression scheme in **Fig. 7** suggests the competing relationship between PKA and calcineurin at excitatory post-synapses. PKA, calcineurin, and PP1 share a post-synaptic locus and enzymatic cascade in the regulation of synaptic plasticity^69–73^. Thus, our results may shed light on the molecular link between sleep regulation and synaptic plasticity control.

## Methods

### Plasmids

Mouse *Prkaca* (NM_008854.5) and *Prkacb* (NM_011100.5) cDNAs were subcloned from mice brain. Mouse *Prakr1a* (NM_021880.4), *Prkar1b.* (NM_001253890.1), *Prkar2a* (NM_008924.3) cDNAs were obtained from pCDNA3-mouse PKA-RIalpha-mEGFP, pcDNA3-mouse PKA-RIbeta-mEGFP, and pCDNA3-mouse PKA-RIIalpha-mEGFP, respectively. pCDNA3-mouse PKA-RIalpha-mEGFP (Addgene plasmid # 45525; http://n2t.net/addgene:45525; RRID:Addgene_45525), pcDNA3-mouse PKA-RIbeta-mEGFP (Addgene plasmid # 45526; http://n2t.net/addgene:45526; RRID:Addgene_45526), and pCDNA3-mouse PKA-RIIalpha-mEGFP (Addgene plasmid # 45527; http://n2t.net/addgene:45527; RRID:Addgene_45527) were gifts from Haining Zhong ^73^. *Prkar2b* cDNA (NM_011158.4) were subcloned from mice brain. cDNAs of Human PP1c (*PPP1CA*, NM_001008709.2; *PPP1CB*, NM_002709.3; *PPP1CC,* NM_002710.4), *PPP1R1A*(NM_006741.4), and PP2Bc (*PPP3CA*, NM_001130691.2; *PPP3CB*, NM_001142353.3) were obtained from GeneScript Japan Inc. (Japan). cDNA with mutations or deletion were constructed by site-directed mutagenesis using PrimeSTAR® HS DNA Polymerase mutagenesis kit (Takara Bio, Japan) following to the manufacturer’s instructions.

For pAAV construction, the cDNA sequence was transferred into the pAAV vector^28^ along with a promoter (*hSyn1*^74^ or *Camk2a*^28^), FLAG tag, Camk2b 3’UTR^28^, WPRE, and SV40 polyA sequences as illustrated in each figure. The mCherry-PKI was constructed based on the AIP2 expression vector^28^ by fusing the PKI 1-24^29^ sequence (TDVETTYADFIASGRTGRRNAIHD for WT; TDVETTYADFIASGATGAANAIHD for 3RA) to the C-terminus of mCherry via a (GGGGS)x3 linker, and assembled in pAAV with the *Camk2a* promoter, dendritic targeting element (DTE) of mouse Map2 gene, WPRE, and SV40 polyA sequences. For double-floxed inverted open reading frame (DIO) constructs, the inverted *Prkaca* sequence flanked by lox2272 and loxP was inserted between the *Camk2a* promoter and the *Camk2b* 3’UTR in the pAAV vector as illustrated in **Extended Data Fig. 4d**. For localized expression, rat PSD95 sequence from FU-dio PSD95-mCherry-W plasmid was fused to the N-terminus of PP1c genes (*PPP1CA*, *PPP1CB*, *PPP1CC*). FU-dio PSD95-mCherry-W was a gift from Elly Nedivi (Addgene plasmid # 73919; http://n2t.net/addgene:73919; RRID:Addgene_73919)^75^ PSD95.FingR sequence were obtained from pCAG_PSD95.FingR-eGFP-CCR5TC and directly fused to N-terminus of *Prkaca* (**Fig. 2**) or *PPP3CA/PPP3CB* (**Fig. 5**). pCAG_PSD95.FingR-eGFP-CCR5TC was a gift from Don Arnold (Addgene plasmid # 46295; http://n2t.net/addgene:46295; RRID:Addgene_46295), and FingR-based constructs used in this study does not include transcriptional control system by the zinc finger-KRAB^36^.

For the construction of pAAV to simultaneously express triple gRNAs, a mCherry expression cassette consisting of *hSyn1* promoter, mCherry, WPRE, hGH polyA sequences was inserted into pAAV, and used as a gRNA template vector. Three sets of U6 promoter and gRNA scaffold containing a gRNA sequence listed in **Extended Data Fig. 5** were tandemly assembled and inserted in upstream of the mCherry expression cassette in the gRNA template vector.

### Animals and sleep phenotyping

All experimental procedures and housing conditions were approved by the Institutional Animal Care and Use Committee of RIKEN Center for Biosystems Dynamics Research and the University of Tokyo. All the animals were cared for and treated humanely in accordance with the Institutional Guidelines for Experiments using Animals. All mice had ad libitum access to food and water and were maintained at ambient temperature and humidity conditions under a 12-hLD cycle. The timing of switching from dark to light environment was set as ZT0. All C57BL/6N mice were purchased from CLEA Japan (Tokyo, Japan). The mice used in each experiment were randomly chosen from colonies. Animal experiments were performed at the University of Tokyo and RIKEN Center for Biosystems Dynamics Research. Triple-CRIPSR KO screening for PKA was conducted at RIKEN and for phosphatases at the University of Tokyo.

### Production of triple-CRISPR KO mice

Guide RNA (gRNA) design and synthesis were performed using the same protocol described in the previous report^76,77^. gRNAs were designed using online design tools including mm10 CRISPR/Cas9 database^23^ (http://www.crispr.riken.jp/), CRISPR-ERA: a comprehensive designer tool for CRISPR genome editing, (gene) repression, and activation (http://crispr-era.stanford.edu/), CRISPRdirect^78^ (http://crispr.dbcls.jp/), and the UNAFold Web Server (http://unafold.rna.albany.edu/). gRNA templates were synthesized with T7 promoter by PCR from the pX330 plasmid (Addegene plasmid # 42230). The T7-fused gRNA templates were amplified by PCR and then used as a template for in vitro transcription with MEGAshortscript T7 kit (Thermo Fisher Scientific, USA). The gRNAs were purified using the MEGAclear kit (Thermo Fisher Scientific, USA). For Cas9 mRNA synthesis, p3s-Cas9HC plasmid^79^ (Addgene plasmid # 43945) was digested with XbaI(New England BioLabs, Japan) and used as a template for in vitro transcription with MEGAshortscript T7 kit (Thermo Fisher Scientific, USA). Cas9 mRNA was purified using the MEGAclear kit (Thermo Fisher Scientific, USA).

One-cell embryo microinjection was performed as described in the previous report^23^. Cas9 mRNA (100 ng/μL) and gRNAs (150 ng/μL in total) were co-injected into the cytoplasm of C57BL/6N fertilized eggs in M2 medium (Merck Millipore, Germany or ARK Resource, Japan) at room temperature (23–25°C). Details of the cytoplasmic injection were reported previously^80^. After microinjection, the embryos were cultured for 1 h in KSOM medium (Merck Millipore, Germany or ARK Resource, Japan) in a 5% CO2 incubator at 37°C, and 25–35 embryos were then transferred to the oviducts of pseudopregnant female ICR mice.

### Genotyping of triple-CRISPR KO mice

Genotyping of KO mice was conducted with the same protocol described in the previous report^23^. Genomic DNA of wild-type and KO mice was prepared from their tails using the DNeasy Blood & Tissue Kit (QIAGEN, Germany), according to the manufacturer’s instructions. qPCR for genotyping was performed using the LightCycler 480 II (Roche, Switzerland) and the SYBR Premix Ex Taq GC (Takara Bio, Japan). Primers for qPCR (**Supplementary Table 1**, Eurofins Genomics, Japan) were annealed to the CRISPR/Cas9 targeting sequences. The absolute target site abundance was calculated using a standard curve obtained from wild-type genomic DNA. The amount of *Tbp*^81^ was quantified and used as an internal control. When the amplified intact DNA by qPCR is less than 0.5% of wild-type genome, we judged that the target DNA is not detectable. When any of three targets was not detected, we classified the animal as a KO. When we could not confirm KO genotype by qPCR, we performed sequencing or a second qPCR using alternative primers, which were independent of first qPCR.

### AAV production and injection

AAV was produced as previously reported^82^ with some modifications. AAV pro 293T (Takara Bio, Japan) cells were cultured in 150 mm dishes (VIOLAMO, AS ONE, Japan) in a culture medium containing DMEM (high glucose) (Thermo Fisher Scientific, USA), 10% fetal bovine serum (FBS) (Biowest, France), and 1% penicillin-streptomycin (PS) (FUJIFILM Wako Pure Chemical, Japan) at 37°C in 5% CO_2_. pAAVs, pUCmini-iCAP-PHP.eB and pHelper plasmid with the plasmid ratio of 1:4:2 based on micrograms of DNA were transfected into cells at >90% confluency with 1mg/ml polyethyleneimine (PEI, Linear, MW 25000, Polysciences). pUCmini-iCAP-PHP.eB for PHP.eB production was a gift from Dr. Viviana Gradinaru (Addgene plasmid # 103005) ^83^. On the next day of transfection, the culture medium was replaced with the culture medium containing DMEM (high glucose, GlutamaxTM, Thermo Fisher Scientific, USA), 2% FBS, 1% MEM Non-Essential Amino Acids solution (NEAA) (Thermo Fisher Scientific, USA), and 1% PS. On the third day following the transfection, the culture medium was collected and replaced with new culture medium also containing DMEM (high glucose, Glutamax^TM^), 2% FBS, 1% MEM NEAA, and 1% PS. The collected culture medium was stored at 4°C. On the fifth day following the transfection, the cells and the culture medium were collected. The suspension of two times collections was separated into supernatant and cell pellet after centrifuge (4000 rpm, 10 minutes). The supernatant mixture with polyethylene glycol solution (PEG) (MW8000, MP Biomedicals, USA) at a final concentration of 8% and left in the ice over 2 hours, whereas AAVs were extracted from the cells pellet which re-suspended in a Tris-MgCl_2_ buffer (10 mM Tris pH 8.0, 2 mM MgCl_2_) followed by 3-4 freeze-thaw cycles in liquid nitrogen. The obtained extract containing AAVs was treated with TurboNuclease (25kU, Accelagen, Australia) for 1 hour. The cells in PEG mixtures were re-suspended by Tris-MgCl_2_ buffer after centrifuge (4000 rpm, 30 minutes) and combined with the lysed cells and incubated in 37°C water for 30 minutes. Then AAVs were purified by ultracentrifugation (OptimaXE-90, Type 70 Ti rotor with 32.4mL OptiSealTM tubes, Beckman Coulter, USA). with 58400 rpm for 145 minutes in 18°C with Iodixanol density gradient solutions (15%, 25%, 40%, and 60% (wt/vol), serumwerk). Viral particles solution was collected and ultrafiltered with Amicon Ultra-15 centrifugal filters (100 kDa, Merck, Germany) to obtain the pure AAVs solution for animal administration.

For AAV titration, virus solution was treated with TurboNuclease in 37°C 1hour followed by Proteinase K (20 mg/ml, 37°C, 1 h). The viral genomes were obtained by phenol: chloroform: isoamyl alcohol 25:24:1 (Nacalai Tesque, Japan) extraction followed by isopropanol precipitation, and were dissolved in 1mmol/L Tris-EDTA (pH 8.0) buffer. The AAVs titer (vg/ml) were quantified depended on the number of WPRE sequences in the AAVs’ genome by qPCR using plasmids as a standard, and WPRE sequences of was amplified by primers (qPCR_WPRE forward:5’-CTGTTGGGCACTGACAATTC-3’; qPCR_WPRE reverse: 5’-GAAGGGACGTAGCAGAAGGA-3’) and qPCR running protocol was 60 s at 95°C for preheating, 45 cycles from 10 s at 95°C to 30s at 60°C using TB Green ® Premix Ex Taq™ GC (Takara Bio, Japan).

For retro orbital injection of AAVs, six-week-old male C57BL/6NJcl mice (CLEA Japan, Japan) were anesthetized with 2%–4% isoflurane inhalation Solution (Pfizer, Japan) and injected with 100 μl of AAVs in their retro orbital sinus. The AAV-administrated mice were subjected to sleep phenotyping at eight-week-old.

### Sleep measurement with SSS

The fully automatic non-invasive respiration-based sleep phenotyping system, called the snappy sleep stager (SSS) was used to monitor the sleep of mice. The SSS recording and analysis were carried out according to the protocol described previously^23^. The light condition of the SSS racks were set to light/dark (12 hours period). In the normal measurement, AAV-administrated 7-week-old C57BL/6N mice or control mice were placed in the SSS chambers for more than one week for sleep recordings, with ad libitum access to food and water. For data analysis, the first day in the chamber was excluded, and analysis was performed for six days of measurement data. For post-natal KO (**Fig. 5**), 8- and 9-weeks old data were analyzed. Sleep parameters such as sleep duration which were defined in previous paper^23^. Sleep staging was performed in every 8-second epoch. The definition of transition probabilities are as follows: *P_WS_* = *N_WS_* / (*N_WS_* + *N_WW_*), and *P_SW_* (transition probability from sleep to wake) is defined as *P_SW_* = *N_SW_* / (*N_SW_* + *N_SS_*), where *N_mn_* is the number of transitions from state m to n (m, n ∈ {sleep, wake}) in the observed period. The balance between *P_WS_* and *P_SW_* determines the total sleep time, i.e., mice with longer sleep time tend to have increased *P_WS_* and/or decreased *P_SW_*. *P_WS_* and *P_SW_* are independent of each other, and it can be deduced from the definition that *P_WS_* + *P_WW_* = 1 and *P_SW_* + *P_SS_* = 1. Amplitude is defined as the coefficient of variation (CV, the standard deviation divided by the mean) of sleep time for each 10-min bin for 24 hours.

### Sleep measurement with EEG/EMG recording

For EEG/EMG recording, mice were implanted with EEG and EMG electrodes for polysomnographic recordings. To monitor EEG signals, 2 stainless steel EEG recording screws with 1.0 mm in diameter and 2.0 mm in length were implanted on the skull of the cortex (anterior, +1.0 mm; right, +1.5 mm from bregma or lambda) and the EEG and EMG electrodes which have 4 pins in 2 mm pitch (Hirose Electric, Japan) with soldered wires were wrapped around the screws. EMG activity was monitored through stainless steel, Teflon-coated wires with 0.33 mm in diameter (AS633, Cooner Wire, USA) connected with electrodes and the other endplaced into the trapezius muscle. Finally, the fixed electrodes were fully covered by dental cement (Unifast III, GC Corporation, Japan) and the mice scalp were sutured. After 7 days of recovery, the mice were placed in experimental cages with a connection of spring supported recording leads. The EEG/EMG signals were amplified (Biotex, Japan), filtered (EEG, 0.5 to 60 Hz; EMG, 5 to 128 Hz), digitized at a sampling rate of 128 Hz, and recorded using VitalRecorder software (KISSEI Comtec, Japan).

For the sleep staging, FASTER method^84^ was used with some modifications to automatically annotate EEG and EMG data. A total of 24 h of recording data were used for the analysis. Sleep staging was performed every 8-s epoch. Finally, the annotations were manually checked. For *Ppp3ca* KO and *Tyr* KO mice (**Fig.5f-i**) and for *Ppp3ca* expression experiments with *Camk2a* promoter (**Fig.6f-h**), three days of automatically annotated data were used for the analyses.

The power spectrum density was calculated for each epoch by fast Fourier transformation (FFT) with Welch’s averaging method. Briefly, each 8-s segment was further divided into 8 overlapping sequences. The overlapping length was 50% of each sequence. The Hamming window was applied onto the sequences before the FFT, and the obtained spectrum was averaged over the 8 sequences. The dirty segments were excluded from the subsequent processes^84^. The power spectrum of each behavioral state (Wake, NREM, and REM) was calculated by averaging the power spectra (1 to 50 Hz) of segments within each state over the observation period. The calculated power spectra were normalized by the total power. The power density in typical frequency domains was calculated as the summation of the powers in each frequency domain (slow, 0.5 to 1 Hz; delta, 0.5 to 4 Hz; theta, 6 to 10 Hz).

Transition probabilities between wakefulness, NREM sleep, and REM sleep were calculated same as previously reported^76^. For example, *P_NW_* = *N_NW_* / (*N_NW_* +*N_NR_* + *N_NN_*), where *N_mn_* is the number of transitions from state m to n (m, n ∈ {wake, NREM sleep, REM sleep}) in the observed period.

### Estimation of EEG delta power dynamics

The simulation and estimation of time constants for the daily increase and decrease of NREM-EEG delta power were carried out as described in the previous study^49^. In brief, Franken and colleagues’ method assumes that the dynamics of EEG delta power in each epoch (S) increases according to Eq (1) during awake and REM sleep phases and decreases according to Eq (2) during NREM sleep.

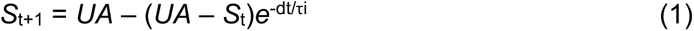

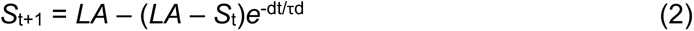

We simulated 72-h EEG recording composed of consecutive 8-s epochs (dt = 8 s). The percentage of delta power for each epoch is used for the calculation and estimation of each parameter. UA is the 99% level of the delta power percentage distribution in the NREM epochs, which is estimated by fitting a gaussian distribution to the histogram of the NREM delta power percentage. LA is defined as the intersection of the distributions of delta power percentage in NREM and REM epochs. τi and τd were estimated by comparing the simulated S and median delta power percentage of sustained (>5 min) NREM episodes. Best τi and τd were explored through the brute force optimization method implemented by the scipy python library^85^.

### ES-mice production

Genetically modified mice were produced using the previously reported ES-mouse method, which allows us to analyze the behavior of F0 generation mice^86,87^. Mouse ES cells (ESCs) were established from blastocysts in 3i medium culture conditions as described previously^88^. Mouse strains used for the ESC establishment were as follows: heterozygous Nex-Cre mice^39^ kindly provided from Carina Hanashima; H11-Cas9, B6J.129(Cg)-*Igs2^tm1.1(CAG-cas9*)Mmw^*/J homozygous mice (The Jakson Laboratory, JAX stock #028239) Male ESCs were cultured as described previously^86,87^. PURECoat amine dishes (Beckton-Dickinson, NJ, USA) was treated with a medium containing LIF plus 6-bromoindirubin-30-oxime (BIO) ^89^ for more than 5 h at 37°C with 5% CO_2_. ESCs were seeded at 1 × 10^5^ cells per well and maintained on the dishes at 37°C in 5% CO_2_ under humidified conditions with a 3i culture medium (Y40010, Takara Bio, Japan) without feeder cells. The ESCs were collected by adding 0.25% trypsin-EDTA solution and prepared as a cell suspension. 10–30 ESCs were injected into each ICR (CLEA Japan, Japan) 8-cell-stage embryo and the embryos were transferred into the uterus of pseudopregnant ICR female mice (SLC, Japan). We determined the contribution of the ESCs in an obtained ES-mouse by its coat color following a previously reported protocol^86,87^. The ES mice uncontaminated with ICR-derived cells were used for the experiment.

### Post-natal knockout by AAV-triple-target CRISPR

6-week-old H11-Cas9 ES mice were administered 2×10^12^ vg of AAVs encoding triple gRNAs downstream of the U6 promoters (AAV-triple-gRNAs). Mice administered the AAVs were subjected to SSS measurements at 8 and 9 weeks old, followed by EEG/EMG recordings.

### Cage change experiment

Cage change experiment was carried out following the six days measurement in SSS. AAV-administered mice were kept in the SSS chambers for three days for baseline recording. On the fourth day, the SSS chambers were replaced with new ones at ZT0 or ZT12 (ZT 0 indicates the beginning of day, or the light phase, and ZT 12 is the beginning of night, or the dark phase). From day 1 of recording, the 6-days data including 3-days baseline were collected and analyzed. The baseline data were averaged and compared with the same time point on the fourth day.

### Statistics

No statistical method was used to predetermine the sample size. The sample sizes were determined based on our previous experiences and reports. Experiments were repeated at least two times with the independent sets of the animals. In the sleep analysis, individuals with abnormal measurement signals or weakened individuals were excluded from the sleep data analyses due to the difficulties in accurate sleep phenotyping.

Statistical analyses were performed by Microsoft Excel and R version 3.5.2. Statistical tests were performed by two-sided. To compare two unpaired samples, the normality was tested using the Shapiro test at a significance level of 0.05. When the normality was not rejected in both groups, the homogeneity of variance was tested using the F-test at a significance level of 0.05. When the null hypothesis of a normal distribution with equal variance for the two groups was not rejected, Student’s t-test was used. When the normality was not rejected but the null hypothesis of equal variance was rejected, a Welch’s t-test was used. Otherwise, a two-sample Wilcoxon test was applied.

To compare more than two samples against an identical sample (e.g., triple-CRISPR screening), the normality was tested with the Kolmogorov-Smirnov test at a significance level of 0.05. When the normality was not rejected in all groups, the homogeneity of variance was tested with Bartlett’s test at a significance level of 0.05. When the null hypothesis of a normal distribution with equal variance was not rejected for all groups, Dunnett’s test was used. Otherwise, Steel’s test was applied.

For multiple comparisons between each group, the Tukey-Kramer test was used when the null hypothesis of a normal distribution with equal variance was not rejected for all groups. Otherwise, Steel-Dwass test was applied.

In this study, p < 0.05 was considered significant (*p < 0.05, **p < 0.01, ***p < 0.001, and n.s. for not significant).

## Acknowledgment

We thank all the lab members at the University of Tokyo and RIKEN Center for Biosystems Dynamics Research (BDR), in particular, Takeyuki Miyawaki, Shiho Sato, Kyoko Shimizu, Yoko Nakano and Kimiko Itayama for AAV preparation; Shuhei S. Sugai for CRISPR experiment; Ayako Shimokawa, Sachiko Tomita, Masako Kunimi, Ruriko Inoue for help with sleep phenotyping; Junko Garcon-Yoshida, Genki N. Kanda, Kazuhiro Kon, Yumika Sugihara, Natsumi Hori, Eriko Matsushita, and Yuichi Uranyu for animal experiment. We also thank members at LARGE, RIKEN BDR for help with ES-mouse production.

This work was supported by grants from the Brain/MINDS JP21dm0207049, Science and Technology Platform Program for Advanced Biological Medicine JP21am0401011, AMED-CREST JP21gm0610006 (AMED/MEXT) (H.R.U.), Grant-in-Aid for Scientific Research (S) JP18H05270 (JSPS KAKENHI) (H.R.U.) and Scientific Research (C) JP23K05738 (JSPS KAKENHI) (K.L.O.), Grant-in-Aid for Early-Career Scientists JP19K16115 (JSPS KAKENHI) (D.T.), HFSP Research Grant Program RGP0019/2018 (HFSP) (H.R.U.), ERATO JPMJER2001 (JST) (H.R.U.) and an intramural Grant-in-Aid from the RIKEN BDR (H.R.U.).

## AUTHOR CONTRIBUTIONS

H.R.U., Y.W., C.S., D.T., and K.L.O. designed the study. Y.W, C.S., D.T., K.L.O., H.F., S.S., K.M. and R.G.Y. performed the sleep phenotype analysis. R.G.Y. performed EEG/EMG data analysis. Y.W., C.S., and D.T. performed AAV production. D.T., R.O., S.S., M.K., M.U.T., H.U., C.H. H.K. and K.S. produced genetically modified mice. H.R.U., Y.W., C.S., D.T., R.G.Y and K.L.O. wrote the manuscript. All authors discussed the results and commented on the manuscript.

## Extended Data

**Extended Data Fig. 1.**
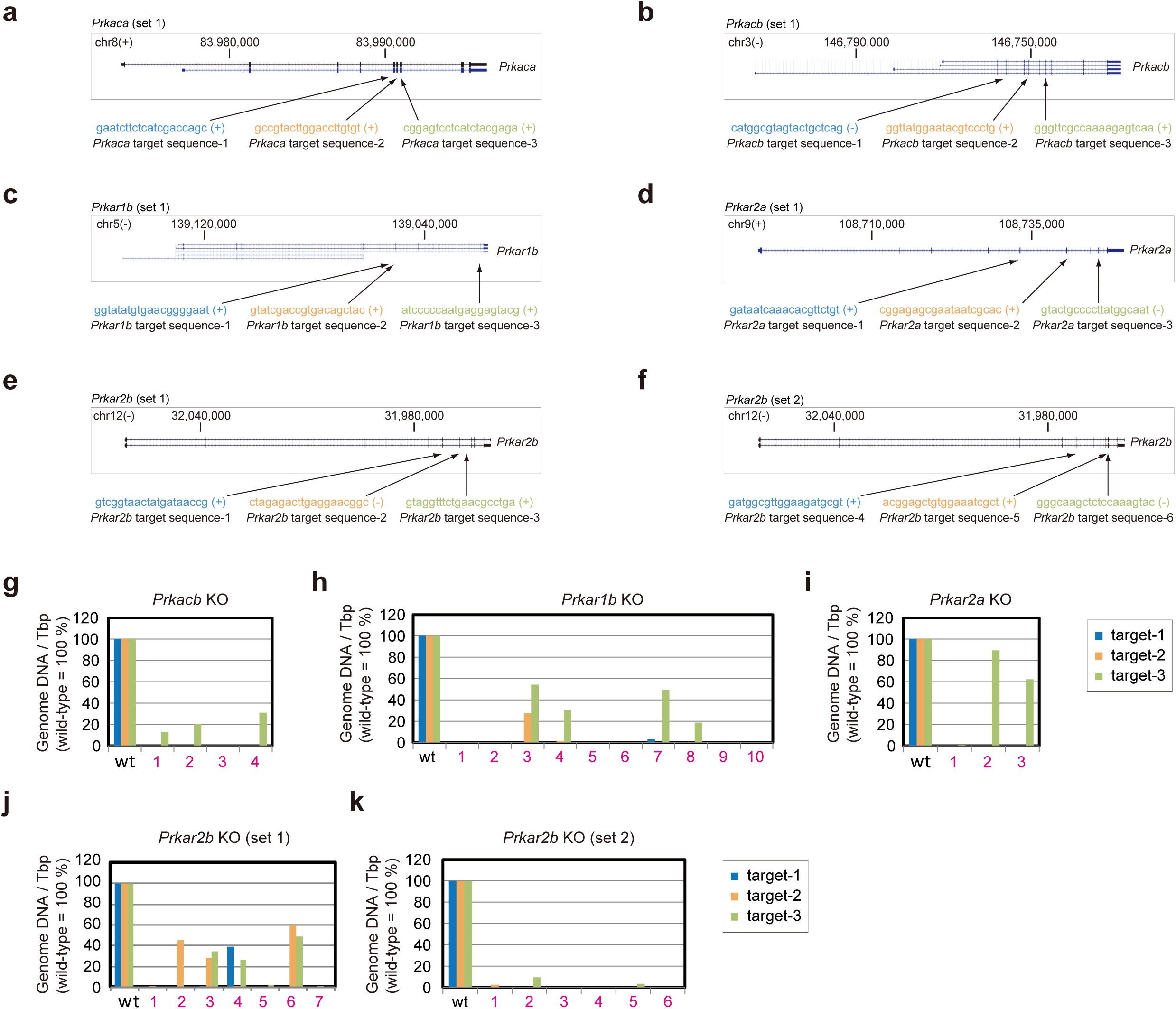
Triple-target CRISPR KO for PKA genes. **(a-f)** Target sequences of the gRNAs for knocking out each *Prkaca*, *Prkacb*, *Prkar1a*, *Prkar1b*, *Prkar2a*, and *Prkar2b* gene. Mouse genomic sequence data were obtained from GRCm39/mm39 via the UCSC Genome Browser (http://genome.ucsc.edu/). The colored letters (blue, orange, and green) show the 20-base target sequences. The target sequences were designed on the sense (+) or the antisense (-) strand of genomic DNA. **(g-k)** The genotyping of *Prkacb* (set1), *Prkar1b* (set1), *Prkar2a* (set1), *Prkar2b* (set1 and set2) KO mice. The qPCR was performed with primer pairs listed in Supplementary Table 1 for the three target sites in a gene. When the 0.5% criteria were met in either set, the mouse was considered a KO mouse. Each number represents each mouse used for the genotyping, and magenta color indicates KO-determined animal. wt, wild-type.

**Extended Data Fig. 2.**
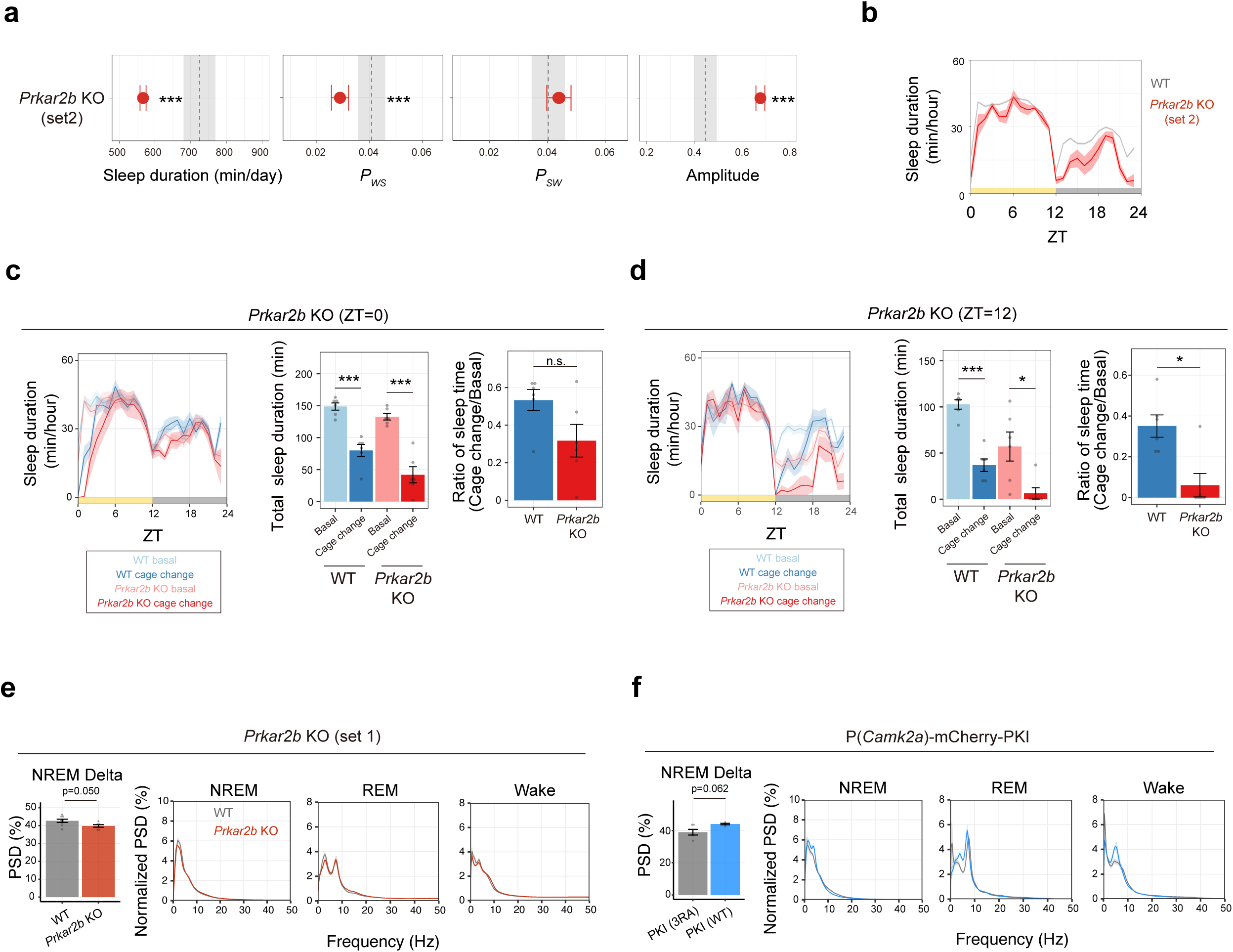
Sleep phenotypes of *Prkar2b* KO and PKI expressed mice. **(a, b)** Sleep/wake parameters (a) and sleep profiles (b) of *Prkar2b* KO (set2) mice (n=6), averaged over 6 days. A gRNA set (set2) independent from **Fig. 1b-f** was used for KO mice production. The black dashed line and the shaded area represent the mean and 1 SD range, respectively, of the wild-type mice wild-type C57BL/6N mice (n=108). Dunnett’s tests were performed between *Prkar2b* KO mice and the wild-type C57BL/6N mice. **(c, d)** Responses of wild-type C57BL/6N mice (n=6) and *Prkar2b* KO mice (n=6) to the cage change stimuli at ZT0 (c) or ZT12 (d). Total sleep duration of 4 hours just after the cage change was used for analysis. “Basal” represents the average of the sleep duration during the same time window over the previous 3 days. “Ratio” represents total sleep duration in cage change response divided by basal. **(e, f)** NREM power density in delta domain (1-4 Hz) and EEG power spectra for *Prkar2b* KO (set2) mice (e) and PKI-expressing mice (f). For *Prkar2b* KO (e), Student’s t-test was performed between wild-type C57BL/6N mice (n=8) and *Prkar2b* KO (set2) mice (n=6), and for PKI-expressing mice (f), Welch’s t-test was performed between PKI (WT) (n=4) and PKI (3RA) mice (n=5). Shaded areas in the line plots represent SEM. Error bars: SEM, *p < 0.05, **p < 0.01, ***p < 0.001. ZT, zeitgeber time; WT, wild-type.

**Extended Data Fig. 3.**
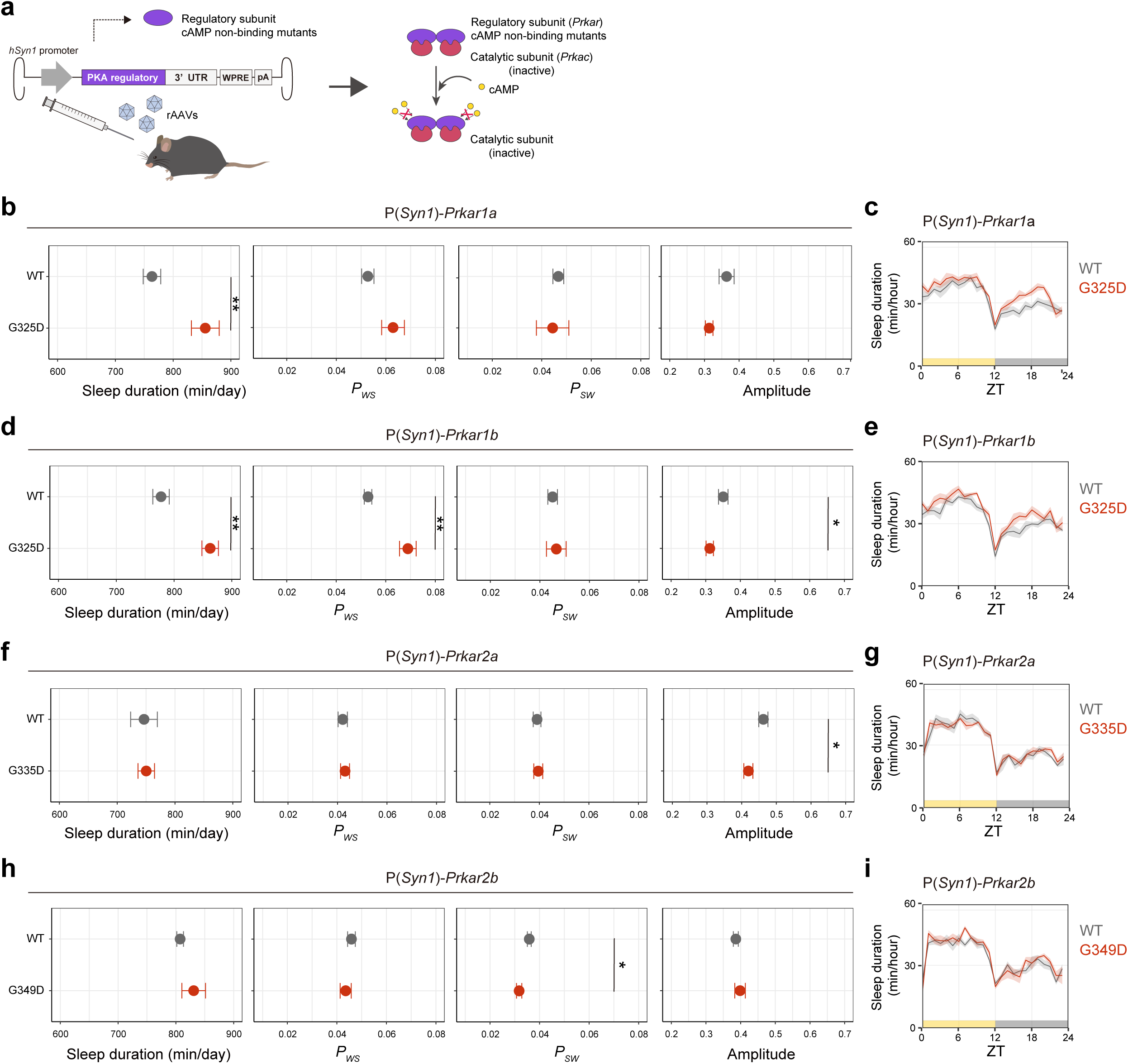
Expression of *Prkar1* and *Prkar2* decreased sleep. **(a)** Schematic diagram of AAV-based PKA regulatory subunit expression. Gly to Asp mutation in the regulatory subunits used in this study block the binding of cAMP to the protein, resulting continued inhibition of the catalytic subunits in the presence of cAMP. **(b, c)** Sleep/wake parameters (b) and sleep profiles (c) of mice expressing *Prkar1a* (WT) (n=6) and its dominant-negative mutant (G325D) (n=6) under the hSyn1 promoter, averaged over 6 days. Welch’s t-test was performed for *P_SW_*, Student’s t-tests were performed for sleep duration, *P_WS_*, and amplitude. **(d, e)** Sleep/wake parameters (d) and sleep profiles (e) of mice expressing *Prkar1b* WT (n=6) and its dominant-negative mutant (G325D) (n=6) under the *hSyn1* promoter, averaged over 6 days. Wilcoxon test was performed for amplitude, and Student’s t-tests were performed for sleep duration, *P_WS_*, and *P_SW_*. **(f, g)** Sleep/wake parameters (f) and sleep profiles (g) of mice expressing *Prkar2a* WT (n=6) and its dominant-negative mutant (G335D) (n=6) under the hSyn1 promoter, averaged over 6 days. Student’s t-tests were performed for comparisons. **(h, i)** Sleep/wake parameters (h) and sleep profiles (i) of mice expressing *Prkar2b* WT (n=6) and its dominant-negative mutant (G349D) (n=6) under the hSyn1 promoter, averaged over 6 days. Welch’s t-tests were performed for sleep duration, Wilcoxon test was performed for amplitude, and Student’s t-tests were performed for sleep duration, *P_WS_*, and *P_SW_*. AAV, adeno-associated virus; *hSyn1*, human synapsin 1; WT, wild-type; ZT, zeitgeber time. Shaded areas in the line plots represent SEM. Error bars: SEM, *p < 0.05, **p < 0.01, ***p < 0.001.

**Extended Data Fig. 4.**
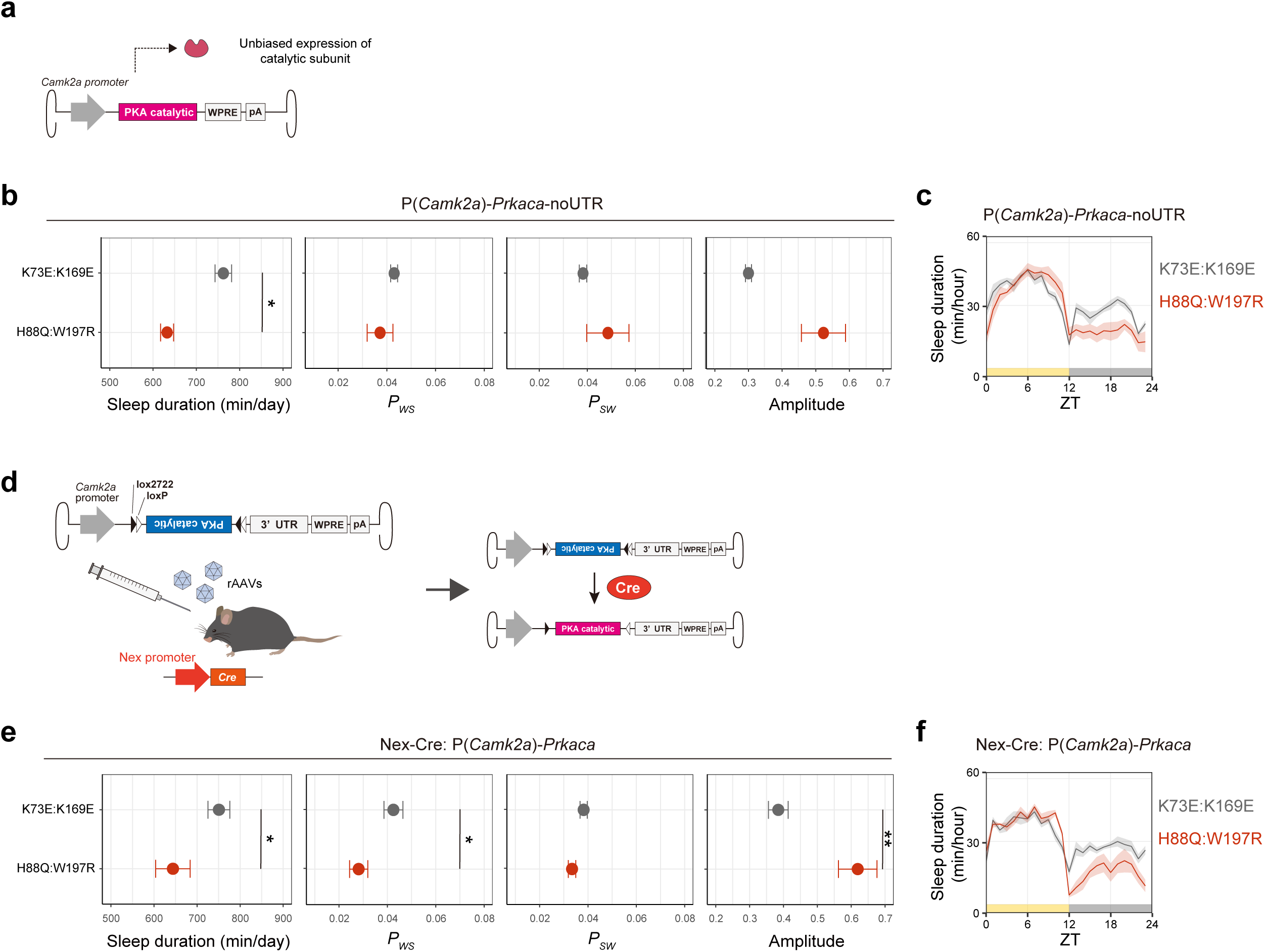
Sleep phenotypes of *Prkaca* expressed mice. **(a)** Schematic diagram of *Prkaca* expression without localized protein fusion. PSD95.FingR was removed from the pAAV construct used in Fig. 2h. **(b, c)** Sleep/wake parameters (b) and sleep profiles (c) of mice expressing *Prkaca* kinase-dead (K73E:K169E) (n=6) and active (H88Q:W197R) (n=6) mutant under the *hSyn1* promoter, averaged over 6 days. Student’s t-tests were performed for the comparisons. **(d)** Schematic diagram of AAV-based PKA catalytic subunit expression in Nex-Cre mice. **(e, f)** Sleep/wake parameters (e) and sleep profiles (f) of Nex-Cre mice expressing *Prkaca* kinase dead mutant (K73E:K169E) (n=6) and active mutant (H88Q:W197R) (n=6) under the *hSyn1* promoter, averaged over 6 days. Wilcoxon test was performed for sleep duration, and Welch’s t-tests were performed for *P_WS_*, *P_SW_*, and amplitude. AAV, adeno-associated virus; WT, wild-type; ZT, zeitgeber time. Shaded areas in the line plots represent SEM. Error bars: SEM, *p < 0.05, **p < 0.01, ***p < 0.001.

**Extended Data Fig. 5.**
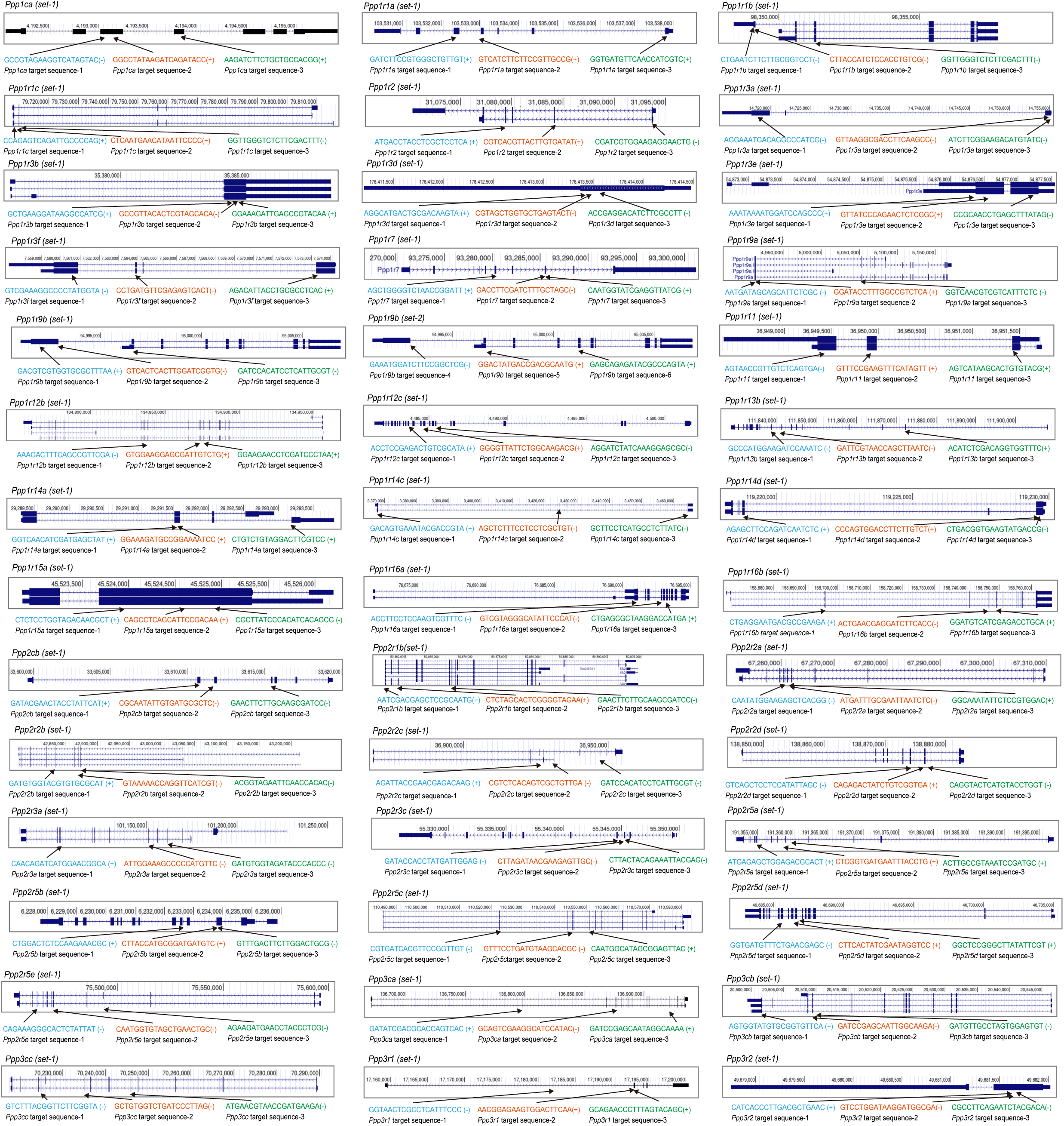
**Triple-target CRISPR KO for PP1, PP2A, and calcineurin genes** Target sequences of the gRNAs for PP1, PP2A and PP2B genes. Each gene had three target sequences. Mouse genomic sequence data were obtained from GRCm39/mm39 via the UCSC Genome Browser (http://genome.ucsc.edu/). The colored letters (blue, orange, and green) show the 20-base target sequences. The target sequences were designed on the sense (+) or the antisense (-) strand of genomic DNA.

**Extended Data Fig. 6.**
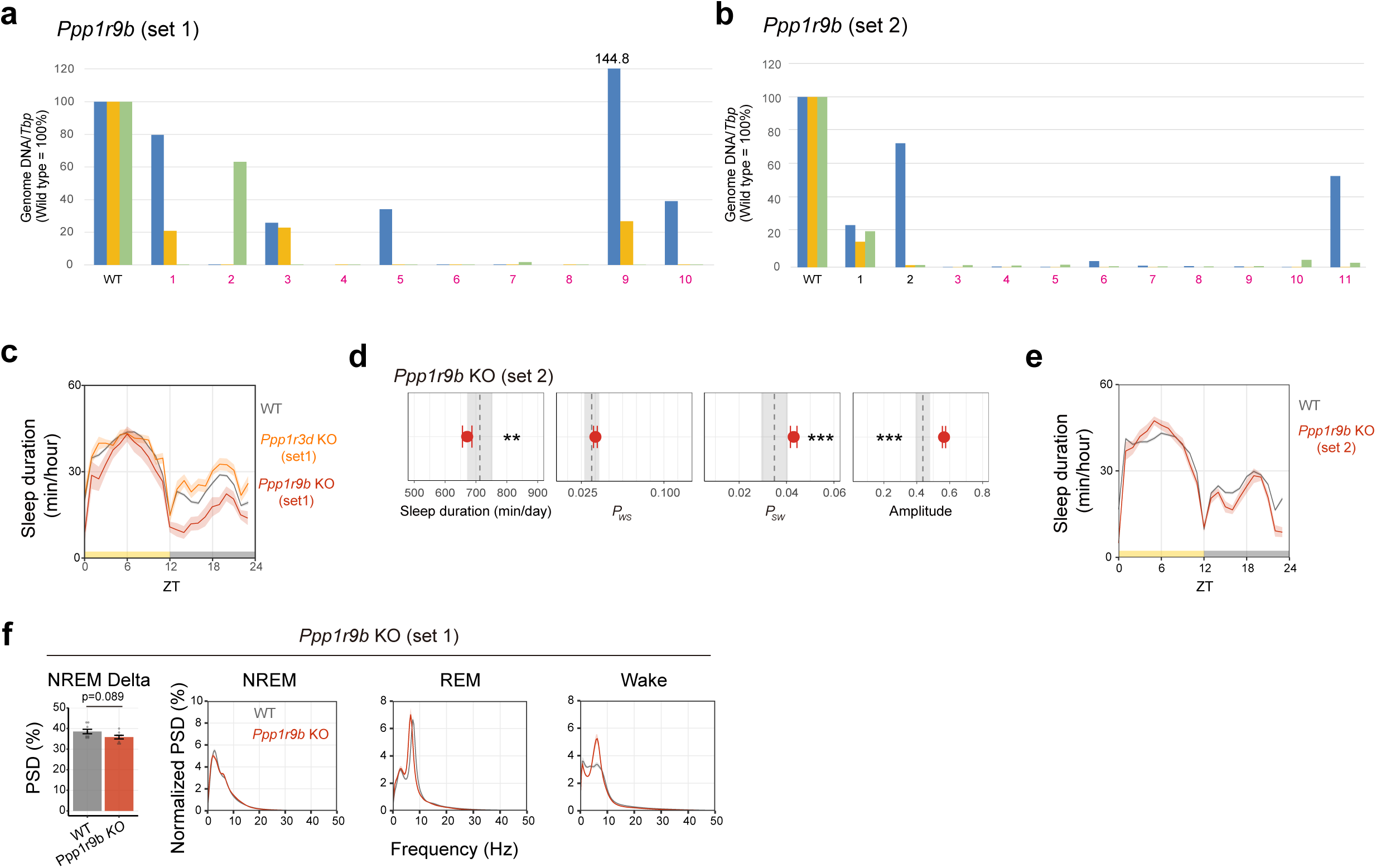
Sleep phenotypes of *Ppp1r9b* KO mice. **(a, b)** The genotyping of *Prkacb* (set1), *Prkar1b* (set1), *Prkar2a* (set1), *Prkar2b* (set1 and set2) KO mice. The qPCR was performed with primer pairs listed in **Supplementary Table 1** for the three target sites in a gene. When the 0.5% criteria were met in either set, the mouse was considered a KO mouse. Each number represents each mouse used for the genotyping, and magenta color indicates KO-determined animal. wt, wild-type. (**c**) 24 hour sleep profiles of *Ppp1r9b* (n=10) and *Ppp1r3d* (n=12) mice KO mice in Figure 3B. **(d, e)** Sleep/wake parameters of *Ppp1r9b* KO (set2) mice (n=9), averaged over 6 days (d) and sleep profiles (e). KO mice were compared with 8-weeks-old wild-type C57BL/6N male mice (n=101). The black dashed line and the shaded area represent the mean and 1 SD range, respectively, of the wild-type mice. Student’s t-tests were performed for comparison. **(f)** NREM power density in delta domain (1-4 Hz) and EEG power spectra of *Ppp1r9b* KO mice (n=8) and wild-type C57BL/6N mice (n=8) shown in Figure 3C-E. Student’s t-test was performed for the comparison in NREM delta power. ZT, zeitgeber time. Shaded areas in the line plots represent SEM. Error bars: SEM, *p < 0.05, **p < 0.01, ***p < 0.001.

**Extended Data Fig. 7.**
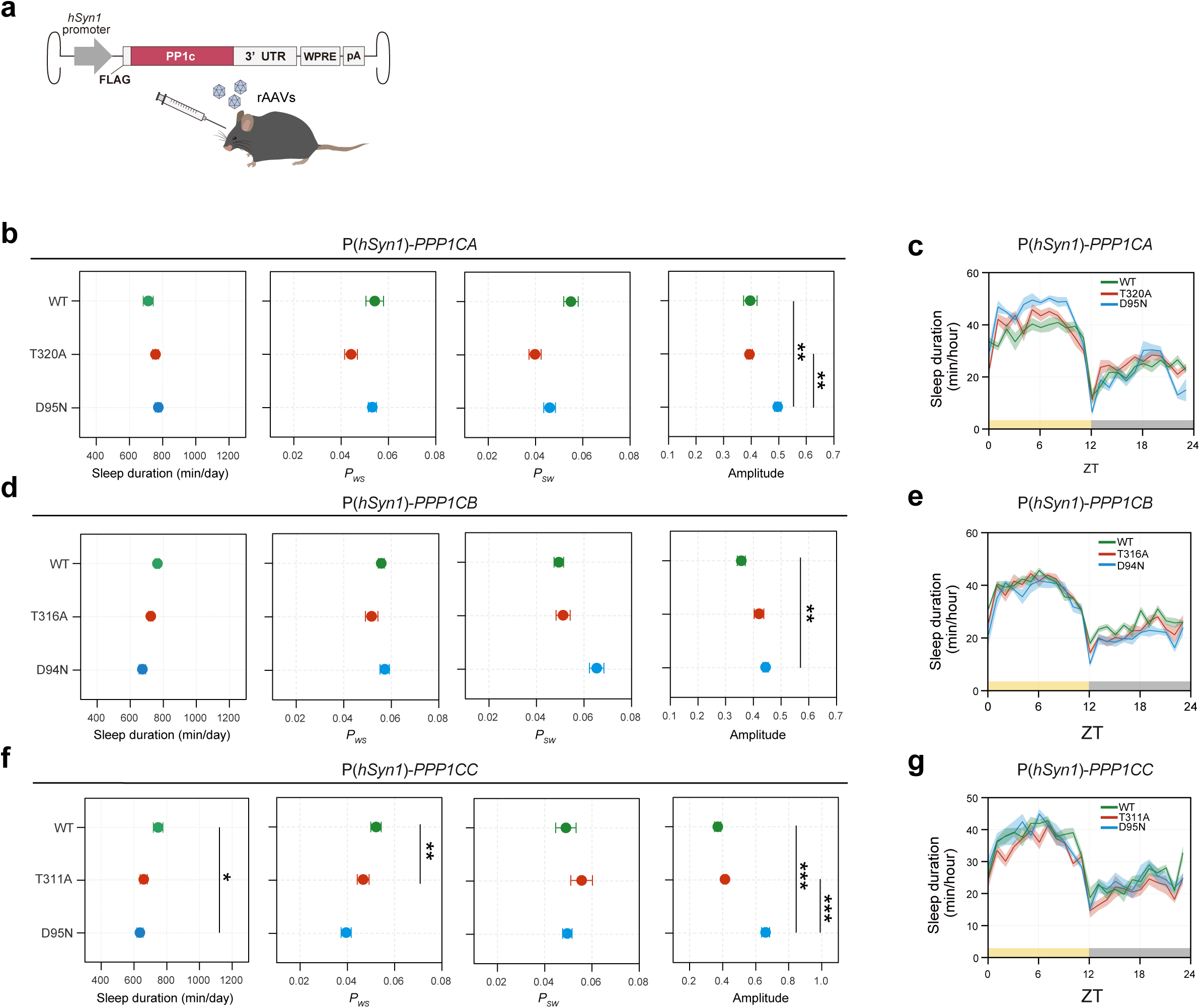
PP1 catalytic subunits without post-synaptic localization do not increase sleep. **(a)** Schematic diagram of AAV-PHP.eB based expression of PP1 catalytic subunits (*PPP1CA*, *PPP1CB*, *PPP1CC*) in mice brain. AAVs were injected into 6-weeks-old male wild-type C57BL/6N mice with the dosage of 4×10^11^ vg/mouse. **(b-g)** Sleep parameters (b, d, f) and 24 hour sleep profiles (c, e, g) of mice expressing PP1 catalytic subunit (n=6, each), averaged over 6 days. The dosages of the AAVs were 4×10^11^ vg/mouse. For *PPP1CA*, *PPP1CB,* and *PPP1CC*, Tukey-Kramer tests were performed for sleep duration, *P_WS_*, *P_SW_* and amplitude between all individual groups. Shaded areas in the line plots represent SEM. Error bars: SEM, *p < 0.05, **p < 0.01, ***p < 0.001. WT, wild-type; ZT, zeitgeber time.

**Extended Data Fig. 8.**
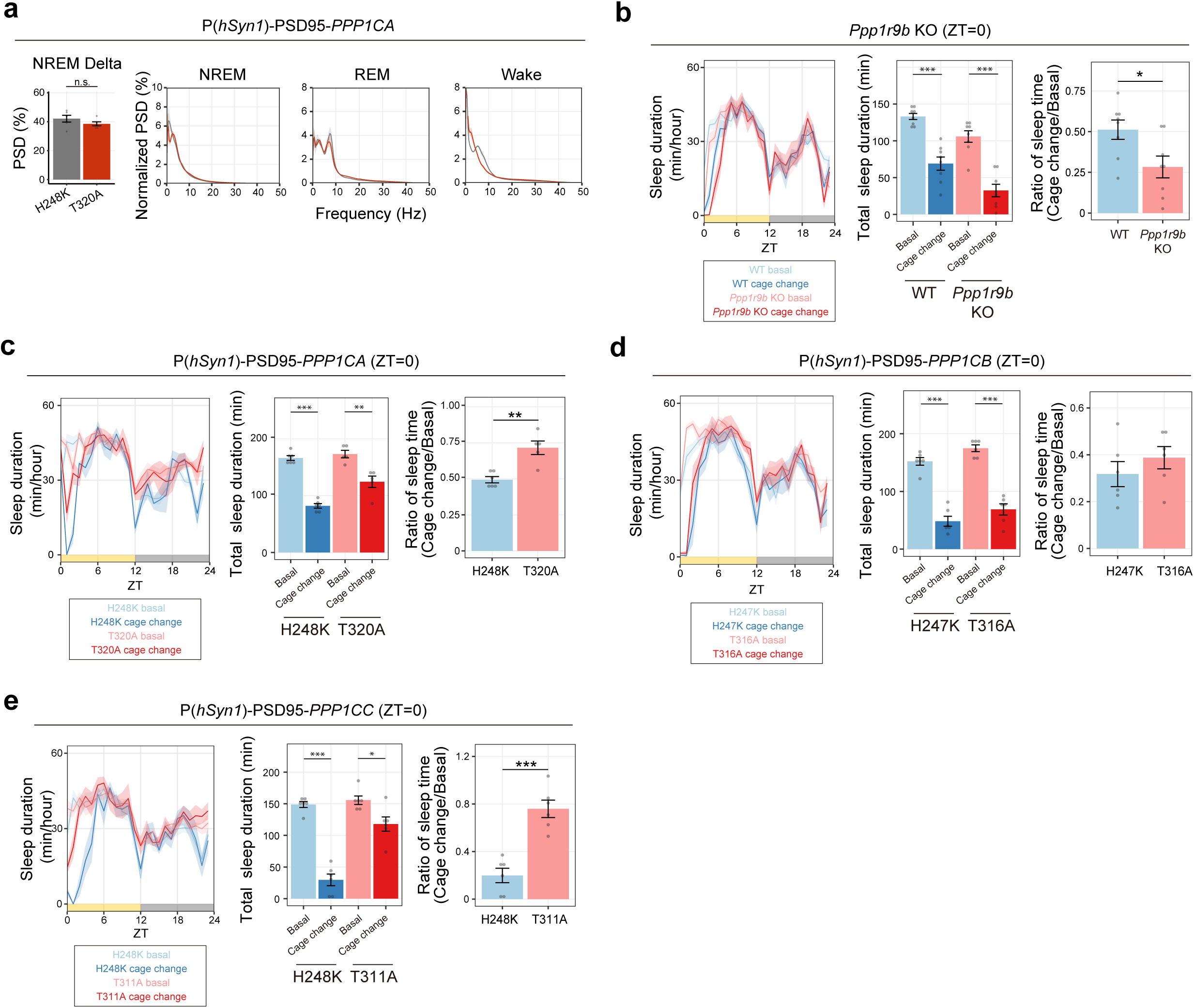
Sleep phenotypes of PP1-expressed mice. **(a)** NREM power density in delta domain (1-4 Hz) and EEG power spectra of *PSD95-fused PPP1CA* phosphatase-dead (H248K) mice (n=5) and its active mutant (T320A) mice (n=5) shown in Figure 4H-J. Student’s t-test was performed for the comparison in NREM delta power. **(b)** Responses of *Ppp1r9b* KO mice (n=8) and wild-type C57BL/6N mice (n=8) mice to the cage change stimuli. Total sleep duration of 4 hours just after the cage change was used for the analysis. “Basal” represents the average of the sleep duration during the same time window over the previous 3 days. “Ratio” represents total sleep duration in cage change response divided by basal. Student’s t-tests were performed for the comparisons. **(c-e)** Responses of mice expressing PSD95-fused PP1 catalytic subunits (n=6, each) to the cage change stimuli. The dosages of the AAVs were 4×10^11^ vg/mouse. Student’s t-tests were performed for the comparisons between phosphatase-dead mutant and active mutant. Shaded areas in the line plots represent SEM. Error bars: SEM, *p < 0.05, **p < 0.01, ***p < 0.001. WT, wild-type; ZT, zeitgeber time.

**Extended Data Fig. 9.**
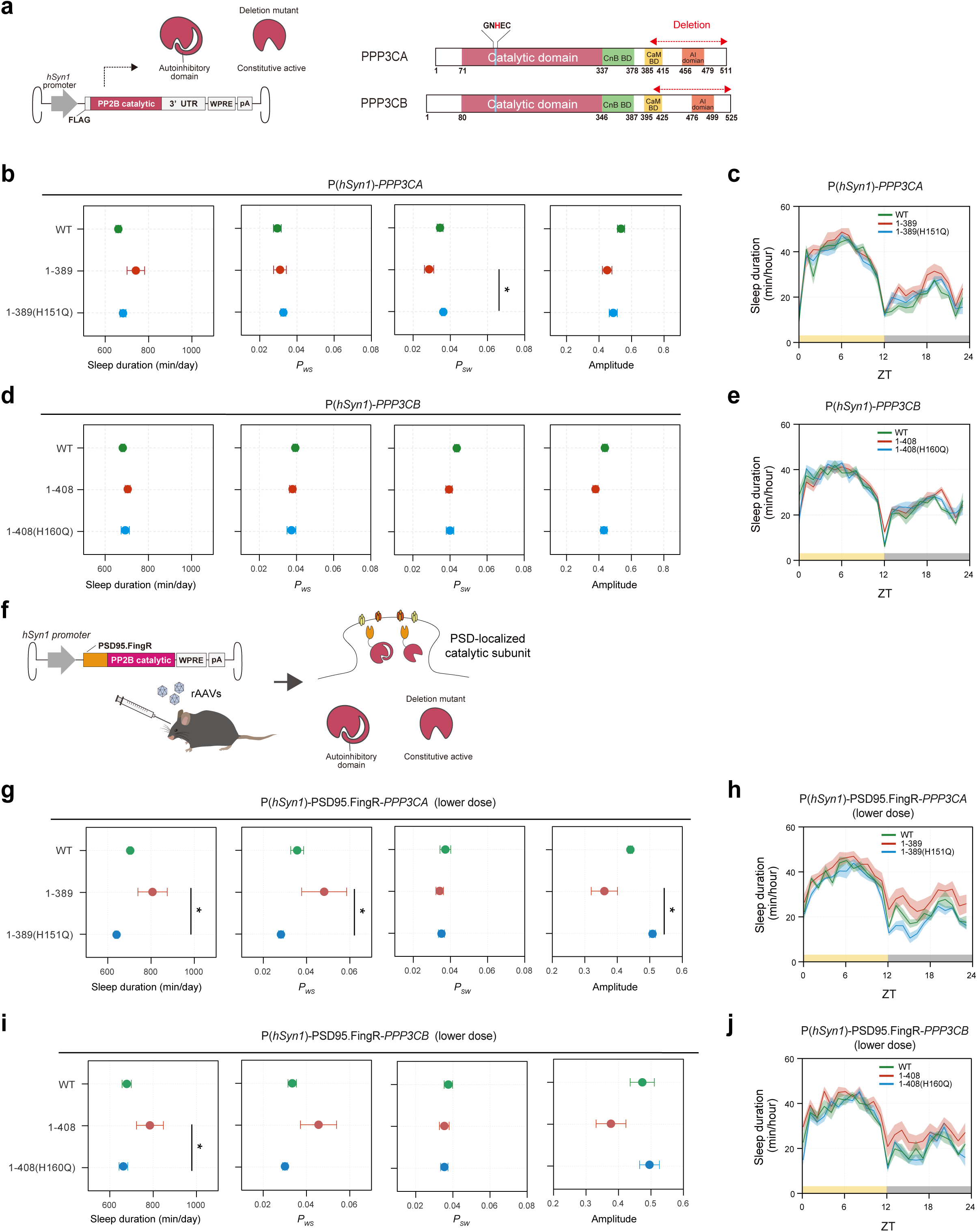
Calcineurin catalytic subunits without post-synaptic localization do not increase sleep. **(a)** Schematic diagram of PP2Bc genes (*PPP3CA*, *PPP3CB*) expression without localized protein fusion. PSD95.FingR was removed from the pAAV construct used in Fig. 5f (left). Information on mutation sites used in this study. Loss of function mutation site was highlighted in red, and the deletion regions for constitutive activation were marked in red dotted line (right). AAVs were injected into 6-weeks-old male wild-type C57BL/6N mice with the dosage of 4×10^11^ vg/mouse. **(b-e)** Sleep parameters (b, d) and 24 hour sleep profiles (c, e) of mice expressing PP2B catalytic subunit (n=6, each), averaged over 6 days. For *PPP3CA*, Steel-Dwass tests were performed for sleep duration, and Tukey-Kramer were performed for *P_WS_* and *P_SW_* and amplitude between all individual groups. For *PPP3CB*, Tukey-Kramer tests were performed for sleep duration, *P_WS_*, *P_SW_* and amplitude between all individual groups. **(f)** Schematic diagram of AAV-PHP.eB based expression of PSD95.FingR-fused PP2B catalytic subunits (*PPP3CA*, *PPP3CB*) in postsynaptic density. The overall scheme is the same as Fig. 5f. The dosages of the AAVs were 2.5×10^10^ vg/mouse (half dosage of Fig. 5f). **(g-j)** Sleep parameters (g, i) and 24 hour sleep profiles (h, j) of mice expressing PSD95.FingR-fused PP2B catalytic subunit (n=6, each), averaged over 6 days. For *PPP3CA*, Tukey-Kramer tests were performed for sleep duration and *P_SW_*, and Steel-Dwass tests were performed for *P_WS_* and amplitude between all individual groups. For *PPP3CB*, Steel-Dwass test was performed for sleep duration and *P_WS_*, and Tukey-Kramer tests were performed for *P_SW_* and amplitude between all individual groups. Shaded areas in the line plots represent SEM. Error bars: SEM, *p < 0.05, **p < 0.01, ***p < 0.001. WT, wild-type; ZT, zeitgeber time.

**Extended Data Fig. 10.**
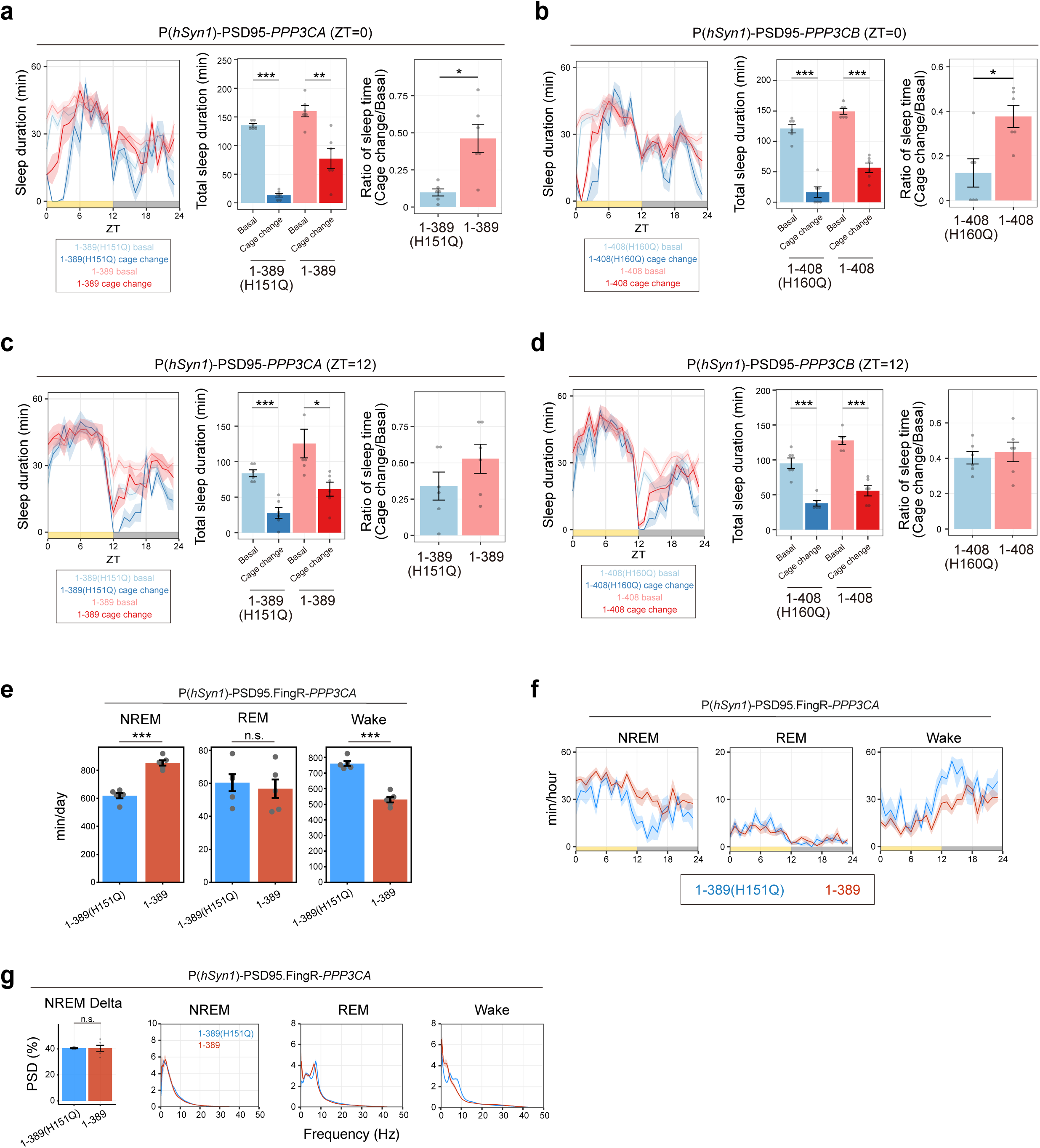
Sleep phenotypes of calcineurin expressed mice. **(a-d)** Responses of mice expressing PSD95.FingR-fused PP2B catalytic subunit (*PPP3CA* and *PPP3CB*) to the cage change stimuli at ZT0 (a, b) or ZT12 (c, d) (n=6 each). Total sleep duration of 4 hours just after the cage change was used for analysis. “Basal” represents the average of the sleep duration during the same time window over the previous 3 days. “Ratio” represents total sleep duration in cage change response divided by basal. Student’s t-tests were performed for the comparisons. **(e, f)** Sleep phenotypes (e) and sleep profiles (f) measured by EEG/EMG recordings for PSD95.FingR-fused *PPP3CA* (1-389) (n=5) and 1-389 (H151Q) (n=5) mutant-expressing mice measured by EEG/EMG. Student’s t-tests were performed for the comparisons. The dosages of the AAVs were 5.0 x 10^10^ vg/mouse. **(g)** NREM power density in delta domain (1-4 Hz) and EEG power spectra of mice expressing *PSD95-fused PPP3CA* inactive mutant 1-389 (H151Q) (n=5) or active mutant (1-389) (n=5). Welch’s t-test was performed for the comparison in NREM delta power. Shaded areas in the line plots represent SEM. Error bars: SEM, *p < 0.05, **p < 0.01, ***p < 0.001. KO, knock out; WT, wild-type; ZT, zeitgeber time.

## Supplementary Table

**Supplementary Table 1** Oligonucleotide sequences used in quantitative PCR (qPCR). The qPCR primers (Eurofins Genomics, Japan) were used for genotyping of knockout mice.

